# B4 Raf-like MAPKKK RAF24 regulates *Arabidopsis thaliana* flowering time through HISTONE MONO-UBIQUITINATION 2

**DOI:** 10.1101/2024.06.12.598286

**Authors:** Qiaomu Li, Le Wang, Lauren E Grubb, Mohana Talasila, Maria Rodriguez, Devang Mehta, Sabine Scandola, RG Uhrig

## Abstract

The timing of flowering is a critical agronomic trait governed by a number of external cues. Despite our genetic understanding of flowering time being well established, we have a limited understanding of how these signals are transmitted to different flowering genes through protein phosphorylation. Here, we characterize a novel B4 Raf-like MAPKKK protein kinase called RAF24, whose mutation results in an early flowering phenotype. Comparative analysis to related B4 Raf-like MAPKKKs indicates that RAF24 uniquely affects flowering time, while phosphoproteome analyses found RAF24 impacts the phosphorylation status of proteins involved in distinct flowering pathways. In particular, we found the RING-type ubiquitin ligase HISTONE MONO-UBIQUITINATION 2 (HUB2) to possess the largest phosphorylation change in *raf24* deficient plants relative to wild-type Arabidopsis and that RAF24 suppresses ligase activity of HUB2 in order to maintain appropriate levels of H2Bub1. Furthermore, we found that RAF24 regulates HUB2 phosphorylation through subclass I and III SUCROSE NON-FERMENTING KINASE 2 (SnRK2) protein kinases; known substrates of B4 RAF-like MAPKKKs. Lastly, using a combination of phospho-mimetic and -ablative plant lines, we validate the importance of HUB2 phosphorylation at S^314^ in regulating flowering time. Collectively, our findings implicate RAF24 as a higher-order flowering regulator, while further implicating HUB2 as a centerpiece of flowering regulation.

## INTRODUCTION

As sessile organisms, land plants are able to perceive and adapt to constantly changing external conditions, such as temperature, light and photoperiod. This ability enables them to adjust their developmental transitions, such as flowering time, in response to diel rhythms or seasonal changes (Mouradov *et al*., 2002; Creux & Harmer, 2019; Bao *et al*., 2020). Premature flowering affects the overall fitness of plants, thereby negatively influencing crop yield and productivity (Gaudinier & Blackman, 2020). To avoid this, plants employ a series of complex regulatory networks. To date, our knowledge of flowering rests largely at the transcriptional level, with numerous transcription factors having been identified to fine-tune flowering time. Ultimately, these networks culminate with *FLOWERING LOCUS C* (FLC), a major flowering repressor, directly regulating the expression of floral integrators, *FLOWERING LOCUS T* (FT) and *SUPPRESSOR OF OVEREXPRESSION OF CONSTANS1* (SOC1), to suppress premature flowering (Lee & Lee, 2010; Deng *et al*., 2011; Bao *et al*., 2020).

Post-translational modifications (PTMs), such as protein phosphorylation and ubiquitination, work synchronously with transcriptional changes to control flowering time (Linden & Callis, 2020). Protein phosphorylation is directed by protein kinases, which transfer the phosphate group from ATP to Ser, Thr and Tyr residues of a protein substrate (Ubersax & Ferrell, 2007), resulting in a multitude of outcomes, including: altered enzymatic activity, protein stability and subcellular localization (Humphrey *et al*., 2015). For instance, calcium-dependent protein kinase CPK28 directly phosphorylates PLANT U-BOX 25 (PUB25) and PUB26 to upregulate its ubiquitin ligase activity and ensure immune homeostasis (Wang *et al*., 2018a). So far, several kinases have been related to flowering time control (Ogiso *et al*., 2010; Chen *et al*., 2020; Sanagi *et al*., 2021; Li *et al*., 2021), including SHAGGY-like kinase 12 (SK12), which was recently resolved to phosphorylate the core flowering regulator CONSTANS (CO) causing rapid degradation of CO by the 26S proteasome, thereby repressing flowering (Chen et al., 2020).

In addition to phosphorylation, protein ubiquitination is also associated with flowering. Protein ubiquitination is mediated by a multi-enzymatic cascade. This involves E1 enzymes, which activate ubiquitin molecules that are then passed to E2 enzymes to form an E2-ubiquitin intermediate. Lastly, E3 enzymes direct the transfer of an ubiquitin moiety from E2 enzymes to specifically recognized protein targets, leading to ubiquitination of substrates (Vierstra, 2009). Interestingly, a number of E2s and E3s have been shown to possess regulatory functions in flowering development. For example, the interaction of RING-type ubiquitin ligases UBC1/2, with HISTONE MONOUBIQUITINATION1 (HUB1) and HUB2 to mono-ubiquitinate histone B2 (H2B) proteins and regulate flowering time (Cao *et al*., 2008; Gu *et al*., 2009). Absence of ubiquitin-modified H2B (H2Bub1) has been suggested to decrease FLC levels leading to early flowering phenotypes in *hub1* and *hub2* mutants (Cao *et al*., 2008; Gu *et al*., 2009). H2Bub1 is also known to modulate the circadian expression of genes (Himanen *et al*., 2012; Woloszynska *et al*., 2019), and to induce light-responsive genes (Bourbousse *et al*., 2012), in addition to modulating the expression of auxin biosynthesis genes (Zhang *et al*., 2021). Most recently, HUB1 and 2 have been shown to interact with SPEN3 and RNA-binding KH domain-containing protein (KHD1), to influence the formation of anti-sense COOLAIR transcripts (Woloszynska *et al*., 2019). Collectively, these findings indicate HUB2 maintains a complex, multifaceted connection to flowering time. Despite HUB2 being extensively linked to multiple biological processes in plants, how HUB2 is modulated in plants remains unknown. Interestingly, the human orthologs of HUB1/2, Ring Finger Protein 20/40 (RNF20/40), are found to be phosphorylated by the ATAXIA-TELANGIECTASIA MUTATED (ATM) protein kinase, which modulates H2Bub1 levels, suggesting that phosphorylation may affect the ubiquitin ligase activity of HUB2 (Moyal *et al*., 2011).

Recently, a subfamily of Arabidopsis B4 Raf-like MAPKKKs (RAF18, RAF20 and RAF24) were reported to phosphorylate sucrose non-fermenting-1 (SNF1)-related kinases (SnRK2s) (MAPK Group, 2002; Lozano-Juste et al., 2020). Upon osmotic stress conditions, RAF18, RAF20 and RAF24 strongly phosphorylate subclass I SnRK2s, which then phosphorylate and activate VARICOSE (VCS) to modulate the mRNA population and plant stress response (Lin et al., 2020; Soma et al., 2020; Fàbregas et al., 2020). In fact, Lin et al also found that RAF24 was able to weakly phosphorylate SnRK2.6, one of the members from subclass III (Lin et al., 2020). On the other hand, some RAFs have been found to be involved in plant growth and fitness. For example, CTR1/RAF1 was characterized as a negative regulator of ethylene hormone signaling (Ju et al., 2012), while EDR1/RAF2 has been implicated in fine-tuning plant immunity (Zhao et al., 2014). In addition, RAF22 and RAF36 were reported to negatively regulate abiotic stress responses and BHP/RAF27 was shown to be a regulator of stomatal opening (Hayashi et al., 2017; Kamiyama et al., 2021; Sun et al., 2022).

Although RAF18, 20 and 24 have been assigned roles in osmotic stress response, other biological roles for these RAFs remain unknown. In this study, we report that RAF24 functions as a flowering time repressor that operates independently of its close relatives; RAF18 and RAF20. Our combined use of genetic, molecular and biochemical analyses coupled with quantitative phosphoproteomics, reveals that RAF24, through select subclass I and III SnRKs, regulates HUB2 to influence flowering time. Moreover, we reveal that the ability of RAF24 to fine-tune flowering seems to be dependent on FLC and FT, but not SOC1. We also find that loss of RAF24 results in changes in both HUB2 ubiquitin ligase activity and its protein interactome. Finally, using a combination of phospho-mimetic (HUB2^S314D^) and -ablative (HUB2^S314A^) mutations, we show that phosphorylation of HUB2 at S^314^ suppresses the early flowering phenotype of the *raf24-2* mutant. Taken together, we reveal RAF24 to be a protein kinase whose function is to repress flowering by influencing HUB2 ligase activity and its interactions with other proteins.

## Results

### Loss of RAF24 causes early flowering in Arabidopsis

The Arabidopsis subgroup B Raf-like MAPKKKs includes seven members. Among these, clade B4 (RAF18, RAF20 and RAF24) are phylogenetically related (Supplemental Figure 1; (MAPK Group, 2002; Lozano-Juste *et al*., 2020)). Previously, RAF18, RAF20 and RAF24 were found to be phosphorylated by subclass I SnRK2 kinases in response to osmotic stress conditions (Stecker *et al*., 2014; Soma *et al*., 2017, 2020; Lin *et al*., 2020). Further, diel phosphoproteomic analyses found RAF18 and RAF24, but not RAF20, to possess a significantly increased phosphorylation status in rosette leaves at the end of day (ED) compared to the end of night (EN), suggesting that B4 clade RAFs may also have additional roles independent of osmotic stress signaling (Supplemental Figure 1) (Uhrig *et al*., 2019). To better understand potential alternative roles for these RAFs, we acquired homozygous T-DNA insertion lines for *RAF18, RAF20* and *RAF24*, which included *raf18-1* (SALK_143032)*, raf20-1* (SALK_053369) and *raf24-2* (SALK_107170), respectively. Here, we found that only *raf24-2* mutants displayed early flowering compared to wild-type *Arabidopsis thaliana* Col-0 (WT Col-0) and related *raf18 / 20* mutants (Figure 1B - C; Supplemental Figure 2A - B). Furthermore, *RAF24* exhibited high transcriptional expression in leaf tissues compared to *RAF18* and *RAF20*; both of which are more abundant in the roots (Soma *et al*., 2020).

**Figure 1.**
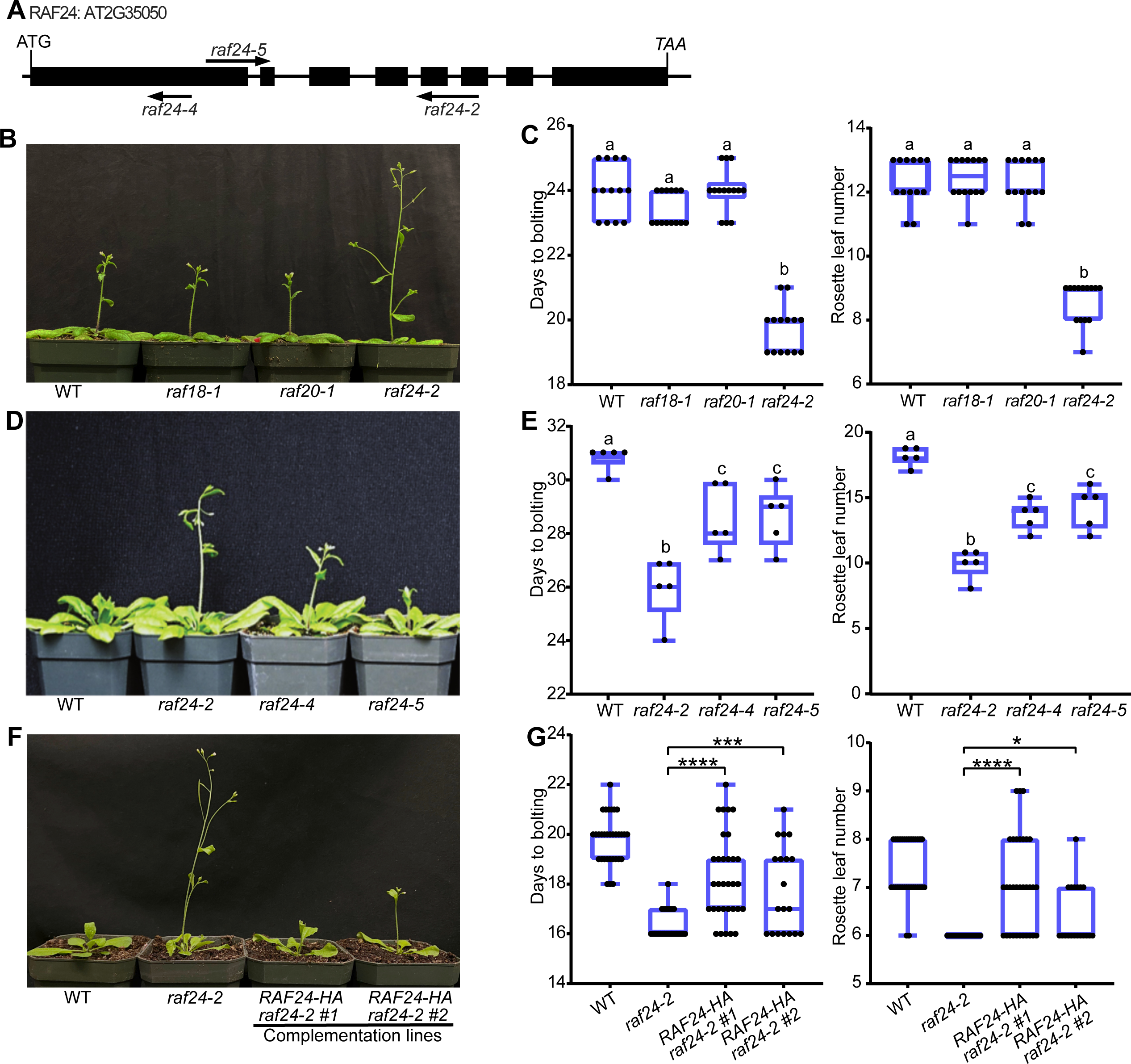
Protein kinase RAF24 regulates flowering time. (A) Schematic genomic diagram of *RAF24* loci. Arrows indicate T-DNA insertion positions; exon 1 of *raf24-4*, exon1 and 2 of *raf24-5* as well as exon 5 and 6 of *raf24-2* mutants. Exons are denoted by black boxes. (B) Early flowering time observed in *raf24* mutants but not in related *raf18* and *raf20* mutants (Supplemental Figure 1). (C) Quantification of flowering time by days to bolting and rosette leaf number at bolting. (D) and (E) Quantification of early flowering in *raf24* mutants by days to bolting and rosette leaf number at bolting. Different letters indicate significant differences among genotypes (one-way ANOVA *p-value < 0.05*). (F) and (G) Flowering phenotype and quantification of WT, *raf24-2* as well as two complementation lines RAF24-HA *raf24-2*. Stars (*) denote one-way ANOVA p-value ≤ 0.05 (*), p-value ≤ 0.001 (***) or p-value ≤ 0.00001 (****).

To further confirm the early flowering phenotype of *raf24-2*, we acquired two additional T-DNA insertional lines, *raf24-4* (GABI_776D02) and *raf24-5* (GABI_702G09) and assessed their time to flowering (Figure 1A). Here, we found *raf24-4* and *raf24-5* loss-of-function lines to also exhibit an early flowering phenotype (Figure 1D - E). Upon comparison, *raf24-2* showed the strongest early flowering phenotype, which corresponded to its lower *RAF24* mRNA levels relative to *raf24-4* and *raf24-5* (Supplemental Figure 2C). In addition to modulating flowering time, we also found that *raf24* mutants have a significantly smaller leaf area compared to WT Col-0 plants, indicating that RAF24 likely has additional developmental roles beyond flowering time regulation (Supplemental Figure 3). Subsequent complementation of the *raf24-2* with *RAF24* reduced time to flowering across multiple lines, confirming the involvement of RAF24 in flowering (Figure 1F – G; Supplemental Figure 2D). Specifically, to assess whether the accelerated flowering phenotype of the *raf24-2* allele is a result of the loss of *RAF24*, we transformed the *RAF24* into the *raf24-2* mutant and quantified the expression of *RAF24* by RT-qPCR (Supplemental Figure 2D). Phenotypic assays showed that two independent *RAF24* complementation lines delayed flowering time relative to *raf24-2*, demonstrating that the observed premature flowering time in *raf24-2* is indeed due to the loss of RAF24 (Figure 1F - G).

### Quantitative phosphoproteomic analysis of *raf24-2* mutant reveals changes in flowering related proteins

To elucidate the underlying molecular mechanisms by which RAF24 modulates flowering, we conducted a quantitative phosphoproteomic analysis of *raf24-2*. Here, we found a decrease in VCS (AT3G13300) phosphorylation, which is consistent with the previously described role of RAF24 in the SnRK2s-VCS module (Soma *et al*., 2017; Kawa *et al*., 2020). We also found several flowering-related proteins to be significantly less phosphorylated in the *raf24-2* mutant compared to WT Col-0 (Table 1; Supplemental Data 1). These included proteins involved in regulating *FLC* expression, such as: VERNALIZATION INDEPENDENCE 4 (VIP4; AT5G61150), ARABIDOPSIS SF1 HOMOLOG (SF1; AT5G51300) and HISTONE MONO-

**Table 1:** Uniquely differentially phosphorylated proteins in *raf24-2*. Quantitative phosphoproteome analysis of *raf18-1*, *raf20-1* and *raf24-2* was performed. Proteins involved in flowering and exhibiting a phosphorylation status decrease unique to *raf24-2* are described. Arabidopsis gene identifier, common name, gene description, predicted localization (SUBA5; https://suba.live/), identified phosphorylation site and corresponding references of research involving each protein are presented.

| Leading Protein | Common name | Description | SUBA5 | Phosphorylation site | Pathway | Reference |
| --- | --- | --- | --- | --- | --- | --- |
| AT3G13300 | VCS | VARICOSE | Cytosol | pSer664 | Osmotic stress signaling | (Soma et al., 2017) |
| AT1G55250 | HUB2 | HISTONE MONO-UBIQUITINATION 2 | Nucleus | pSer314 | Flowering pathway | (Cao et al., 2008; Woloszynska et al., 2019; Gu et al., 2009) |
| AT5G61150 | VIP4 | VERNALIZATION INDEPENDENCE 4 | Nucleus | pSer600<br>pSer605 | Autonomous flowering pathway | (Zhang and van Nocker, 2002) |
| AT5G51300 | SF1 | SPLICING FACTOR 1 | Nucleus | pSer77 | Thermosensory flowering pathway | (Lee *et al.*, 2020) |
| AT1G51140 | FBH3 | FLOWERING BHLH 3 | Nucleus | pSer259 | Photoperiodic flowering pathway; Osmotic stress signaling | (Ito *et al.*, 2012) |
| AT1G08420 | BSL2 | BRI1 SUPPRESSOR 1 (BSU1)-LIKE 2 | Nucleus | pSer973 | BR signaling | (Kim et al., 2016) |
| AT4G03080 | BSL1 | BRI1 SUPPRESSOR 1 (BSU1)-LIKE 1 | Nucleus | pSer491 | BR signaling | (Kim et al., 2016; Kim et al., 2018) |
| AT1G19350 | BES1 | BRI1-EMS-SUPPRESSOR 1 | Nucleus | pTyr175 | Photoperiodic flowering pathway; BR signaling | (Wang *et al.*, 2019) |
| AT1G78700 | BEH4 | BES1/BZR1 HOMOLOG 4 | Nucleus | pSer310<br>pSer312 | BR signaling | (Galstyan and Nemhauser, 2019) |

subcellular localization
pSer = phosphorylated serine
pTyr = phosphorylated tyrosine

UBIQUITINATION 2 (HUB2) (Zhang & van Nocker, 2002; Lee *et al*., 2020). In addition, several FLC-independent flowering phosphoproteins also exhibited a decreased phosphorylation status in *raf24-2* plants (Table 1). These included: BRI1-EMS-SUPPRESSOR 1 (BES1; AT1G19350), a positive regulator of photoperiodic flowering through the promotion of FT expression and FLOWERING BHLH 3 (FBH3; AT1G51140), a flowering regulator that promotes the expression of CO (Wang *et al*., 2019; Sanagi *et al*., 2021). Taken together, this suggests that RAF24 may have wide-ranging, higher-order control over flowering signaling. Among these flowering-related phosphoproteins, phosphorylation of HUB2 (AT1G55250) at site S^314^ exhibited the most dramatic decrease in abundance, being below detection in *raf24-2* relative to WT Col-0 (Table 1). Consequently, we hypothesized that RAF24 plays a key role in regulating flowering time through modulation of HUB2 phosphorylation at S^314^.

### RAF24 and HUB2 co-localize to the nucleus

To better understand the molecular role of RAF24, we next investigated the subcellular localization of RAF24 through the transient expression of c-terminal YFP-tagged RAF24 (RAF24-YFP) in *Nicotiana benthamiana* leaf epidermal cells. Consistent with previous observations (Soma *et al*., 2020), RAF24-YFP is predominantly localized to the cytoplasm (Supplemental Figure 4), with smaller quantities of RAF24-YFP found to localize to the nuclei (Supplemental Figure 4). Next, we then transiently co-expressed RAF24-YFP and mCherry-HUB2 in the *N. benthamiana* to test if they co-localize in the plant cell. Here, we see that a major proportion of RAF24-YFP co-localizes with mCherry-HUB2 in the cell (Figure 2).

**Figure 2.**
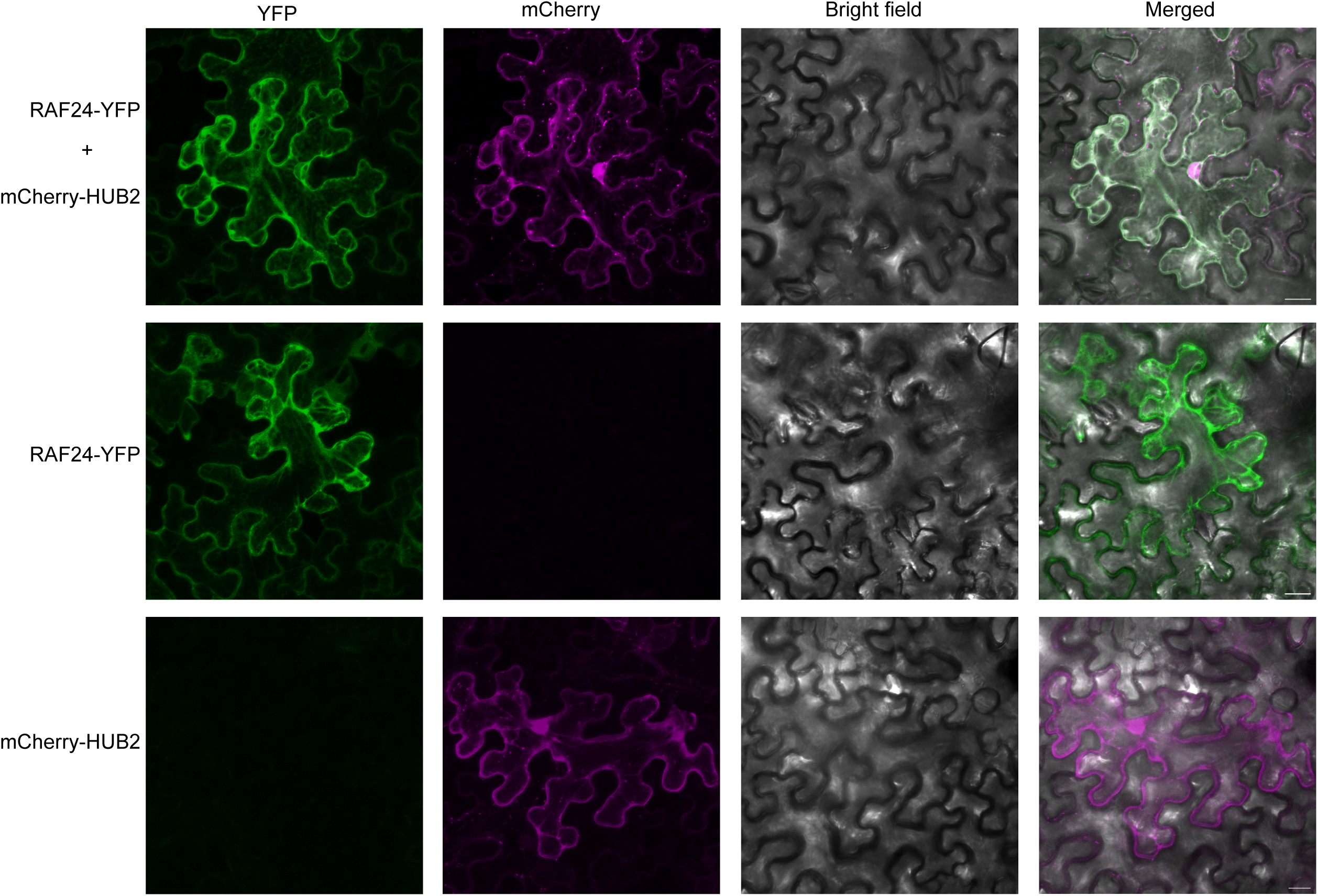
RAF24 co-localizes with HUB2. RAF24-YFP and mCherry-HUB2 were transiently expressed in *N. benthamiana* leaves (See materials and methods). Fluorescence signals were visualized by confocal microscopy 2 days after injection. Scale bars = 20 μm.

### Genetic analysis reveals RAF24 and HUB2 control flowering time and the expression of *FLC* and *FT*

As a result of previous studies demonstrating *hub2* deficient plants to possess an early flowering phenotype (Cao *et al*., 2008; Gu *et al*., 2009), we next explored the potential of a genetic interaction between RAF24 and HUB2 by generating double loss-of-function plant lines. Under a 16:8 photoperiod, *raf24-2* and *hub2-1* single mutants flower earlier than Col-0 as expected, while *raf24-2 hub2-1* double mutants were found to flower slightly earlier than *hub2-1*, but not *raf24-2* (Figure 3A - B). However, upon evaluating the same mutant lines under a 12:12 photoperiod, we found that the *raf24-2 hub2-1* plants flower dramatically earlier than both parental single mutants (Figure 3C - D). Collectively, this indicates that RAF24 and HUB2 act synergistically to influence flowering time. The additive effects of *raf24-2 hub2-1* in accelerating flowering time also indicates likely intersections with the other flowering targets observed to possess reduced phosphorylation in the *raf24-2* mutant.

**Figure 3.**
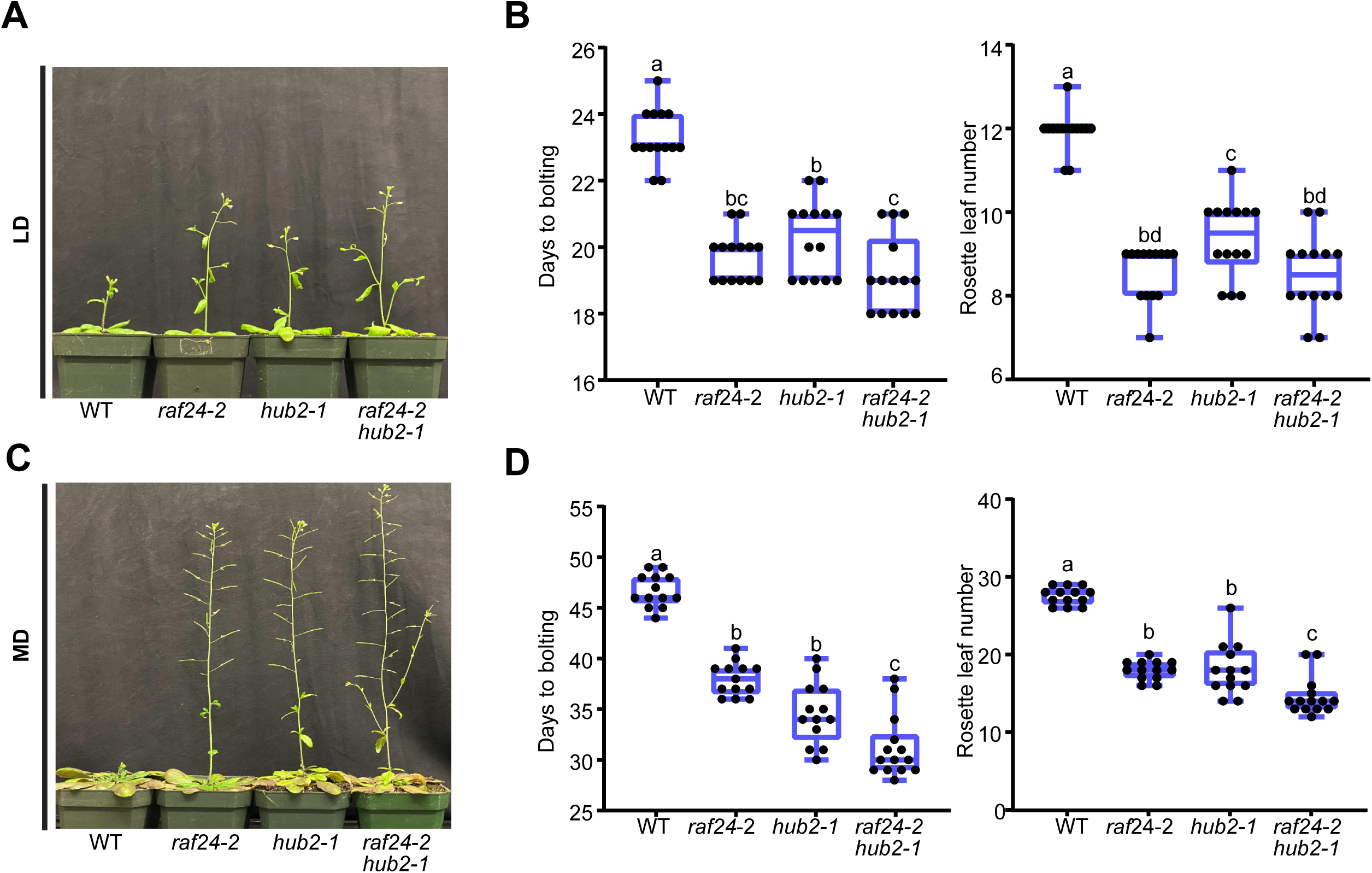
Genetic interaction analysis of HUB2 and RAF24. (A) and (B) Flowering time comparison of *raf24-2*, *hub2-1* and *raf24-2 hub2-1* double mutants relative to WT. 29-day-old plants were photographed under 16:8 photoperiodic conditions and flowering initiation was scored by days to bolting and rosette leaf number at bolting. Calculations were conducted based on more than 12 plants of each genotype. (C) and (D) Flowering time comparison of *raf24-2*, *hub2-1* and *raf24-2 hub2-1* double mutants relative to WT. 56-day-old plants were photographed after being grown under a 12:12 photoperiod. Flowering time was scored by days to bolting and rosette leaf number at bolting. Letters indicate significant differences between genotypes (one-way ANOVA *p-value < 0.05*).

HUB2 is known to regulate flowering pathways by modulating *FLC* transcription through the mono-ubiquitination of H2B (Cao *et al*., 2008; Gu *et al*., 2009). Therefore, we hypothesize that RAF24 may also affect FLC mRNA levels by modulating HUB2 phosphorylation to fine-tune flowering time. To assess this, we examined *FLC* expression levels in *raf24-2*. Consistent with the accelerated flowering time phenotype, *FLC* transcription is significantly decreased in *raf24-2* mutants relative to WT Col-0 (Figure 4A). Simultaneously, we also assessed the expression levels of *FT* and *SOC1*, downstream floral integrators whose expression levels are directly targeted by FLC. Here, we find that *FT* expression is significantly higher, while *SOC1* expression remains unchanged, indicating that RAF24 regulates floral development via FLC and FT (Figure 4A). We next validated that flowering time repression by RAF24 is FLC and FT dependent by crossing *raf24-2* with *flc-6* and *ft-10* mutants, respectively. Here, *raf24-2 flc-6* double mutants flowered just as early as each single mutant plant, respectively, suggesting that the loss of *FLC* was epistatic to *RAF24* and the premature flower development in *raf24-2* was due to *FLC* function (Figure 4B & C). Conversely, *raf24-2 ft-10* plants flowered an average of 7.2 days earlier than *ft-10.* Collectively, this suggests that RAF24 modulates floral development in a FLC and FT dependent manner (Figure 4B & C).

**Figure 4.**
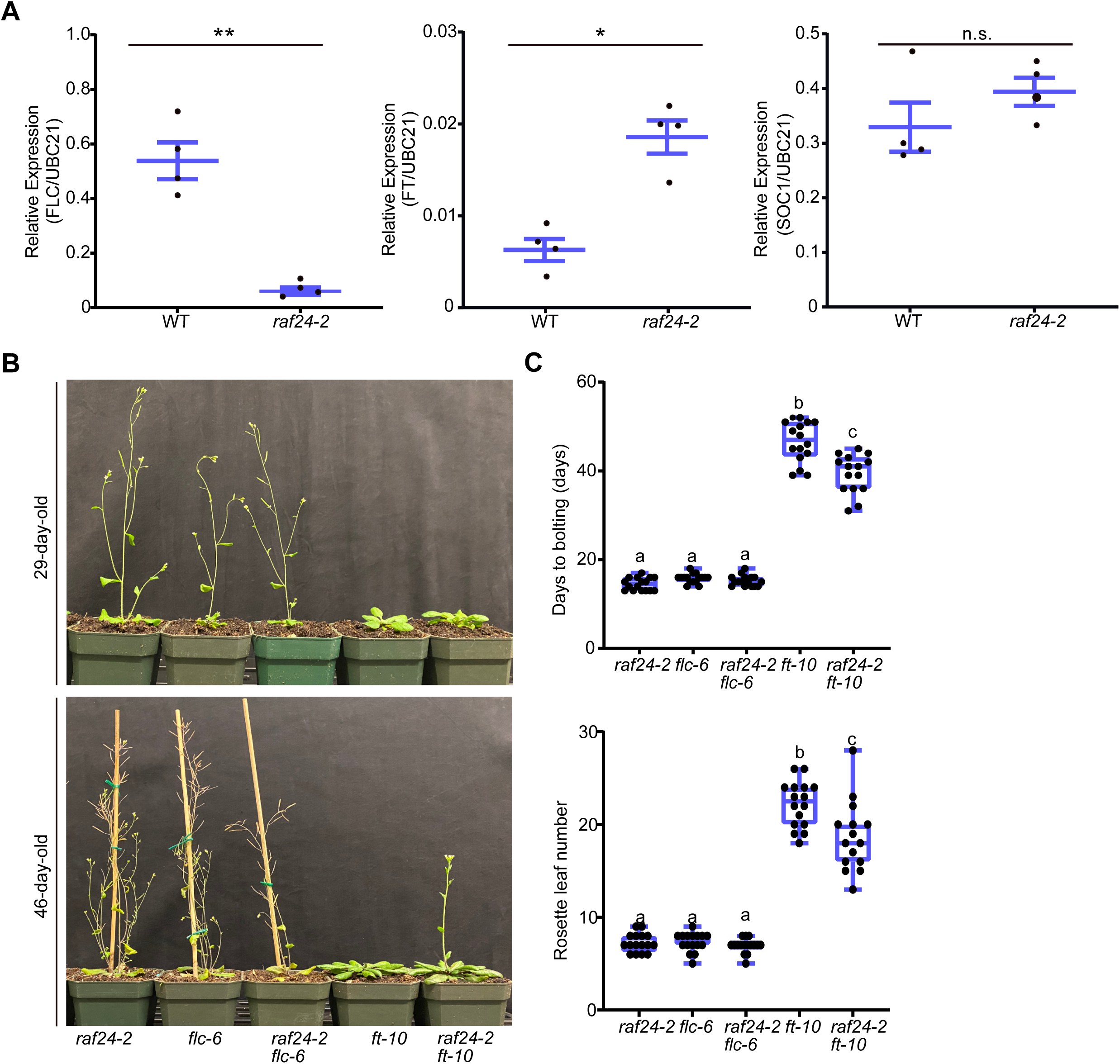
RAF24 fine-tunes flowering time through FLC and FT. (A) Transcriptional expression of *FLC*, *FT* and *SOC1* in 14d old WT and *raf24-2* plants. Stars denote a Student’s t-test *p-value* ≤ *0.05* (*) and *p-value* ≤ *0.01* (**). (B) and (C) Flowering time comparison of *raf24-2 flc-6* and *raf24-2 ft-10* double mutants. 29-day-old plants (upper) and 46-day-old plants (bottom) were photographed (B), with flowering time determined by days to bolting (upper) and rosette number at bolting (bottom) (C). Letters indicate significant differences between genotypes (one-way ANOVA *p-value < 0.05*).

### RAF24 suppresses mono-ubiquitination of H2B by HUB2

HUB2 is responsible for H2B mono-ubiquitination, which has been suggested to promote expression of flowering genes (Cao *et al*., 2008; Gu *et al*., 2009). In humans, the phosphorylation of RNF20/40 (the human homolog of HUB1/HUB2) (Moyal *et al*., 2011) promotes H2Bub1 and activation of DNA repair responses. Therefore, we hypothesized that RAF24 may regulate the phosphorylation status of HUB2 in order to modulate H2Bub1 levels, thereby influencing FLC expression and the control of flowering time. To test this, we performed immunoblotted WT Col-0 and *raf24-2* plants constitutively over-expressing HUB2 using anti-H2Bub1 and -H2B antibodies (Figure 5A & B). Here, we found a higher accumulation of H2Bub1, but not HUB2, in *raf24-2* plants over-expressing a tandem affinity purification (TAP)-tagged HUB2 construct (TAP-HUB2), suggesting that RAF24 suppresses H2B mono-ubiquitination by phosphorylating HUB2 (Figure 5A, B & C). At the same time, we find that *raf24-2 hub2-1* and *hub2-1* plants maintain no H2Bub1 as expected, validating the effectiveness of the anti-H2Bub1 antibodies (Supplemental Figure 5). Surprisingly, we find that TAP-HUB2/Col-0 plants also display an early flowering phenotype when compared to WT Col-0 as previously reported (Woloszynska *et al*., 2019) (Figure 5D - E), suggesting HUB2 likely operates as an integration point for multiple flowering related pathways, with optimal HUB2 levels required for fine-tuning flowering time. Moreover, over-expression of HUB2 in *raf24-2* induces earlier flowering than in WT Col-0 (Figure 5D - E), implying that accumulation of H2Bub1 may also trigger inappropriate flowering time. Collectively, this evidence indicates that RAF24 operates upstream of HUB2 to suppress H2Bub1 in order to fine-tune FLC expression and appropriately regulate flowering time.

**Figure 5.**
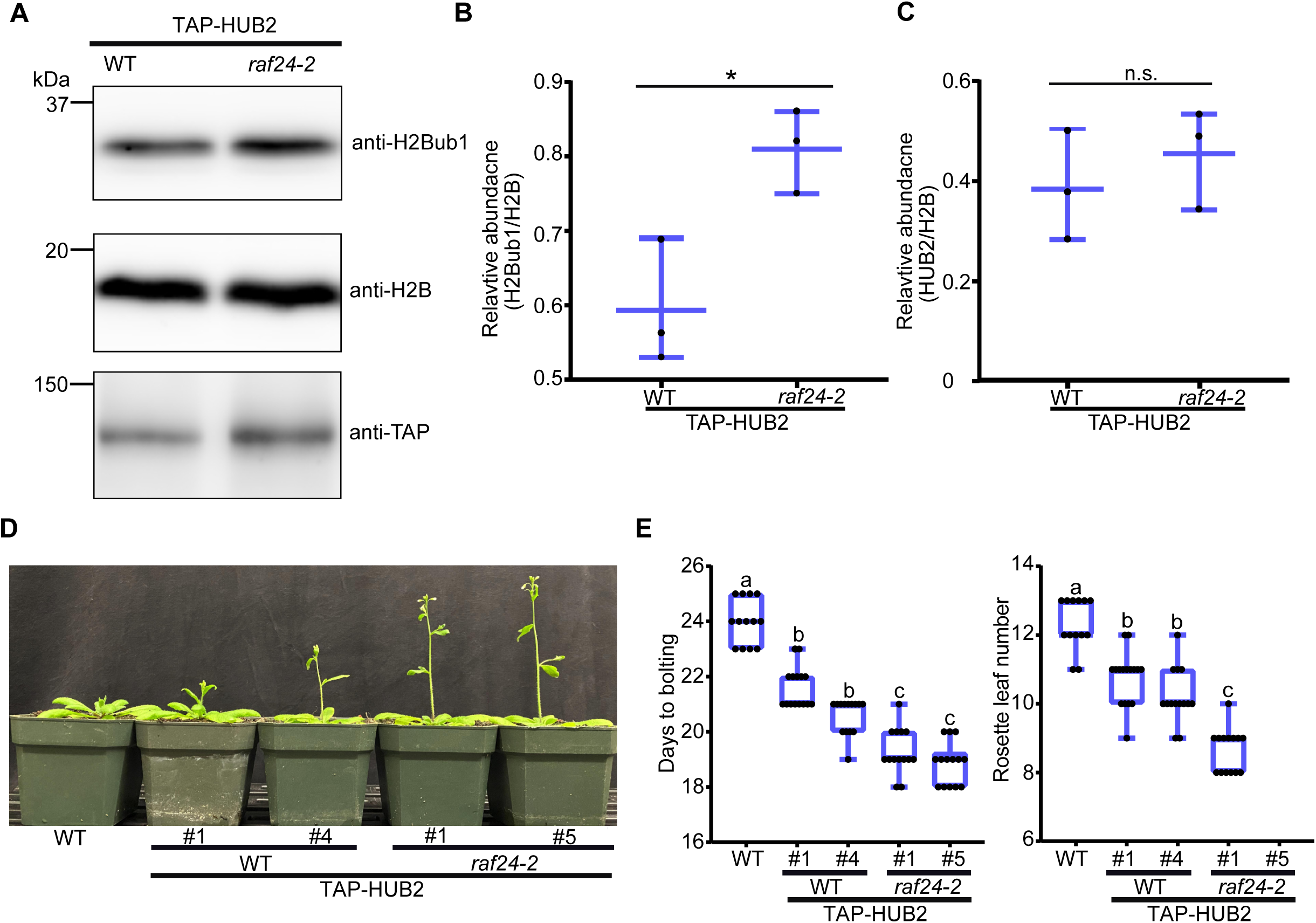
RAF24 suppress the mono-ubiquitination of H2B by HUB2. (A-C) Loss of RAF24 increases H2Bub1 levels. Western immunoblot analysis of 10d-old WT and *raf24-2* plants constitutively over-expressing TAP-HUB2. 50 μg of protein samples were loaded in each lane. Biological replicates (n=3) were examined and quantified with ImageJ software, with levels of H2Bub1 normalized to H2B levels (B) or levels of TAP-HUB2 normalized to H2B levels (C). (D-E) Over-expression of TAP-HUB2 in *raf24-2* also leads to an early flowering phenotype. Two independent lines of 29d-old WT and *raf24-2* plants (denoted by the number) constitutively expressing TAP-HUB2 were grown and imaged (D), with flowering time determined by days to bolting (left) and rosette number at bolting (right) (E). Letters indicate significant differences between genotypes (one-way ANOVA *p-value < 0.05*).

### RAF24 affects protein-protein binding of HUB2

By using RT-qPCR assays, we found that loss of *RAF24* does not affect the expression of either HUB1 or HUB2 (Supplemental Figure 6; Supplemental Data 2), suggesting that RAF24 impacts HUB2 at the protein level, potentially through protein-protein interactions. As previously reported, mono-ubiquitination of H2B is suggested to rely on interactions between UBC1/2 and HUB1/2 as well as the binding between HUB2 and H2B (Liu *et al*., 2007; Cao *et al*., 2008). To better understand this, we performed affinity purification mass spectrometry (AP-MS) experimentation using TAP-HUB2 constructs expressed in either WT Col-0 or *raf24-2* (Supplemental Data 3). Here, we quantified all known interactors of HUB2, except for UBC1/2, in both backgrounds, however, a direct interaction between HUB2 and UBC1/2 has only been shown using yeast 2-hybrid and has not been validated *in vivo*. Importantly, we find the interaction between TAP-HUB2 and HUB1 as unchanging when quantified in the presence or absence of RAF24, suggesting that the phosphorylation of HUB2 is not vital for heterodimer formation between HUB1 and HUB2 (Figure 6A). However, TAP-HUB2 is able to immunoprecipitate significantly more H2B (AT5G22880) in *raf24-2* plants than that of WT Col-0, implying that RAF24-induced phosphorylation of HUB2 dampens its affinity for H2B as a substrate, suggesting a mechanism by which RAF24 suppresses H2Bub1.

**Figure 6.**
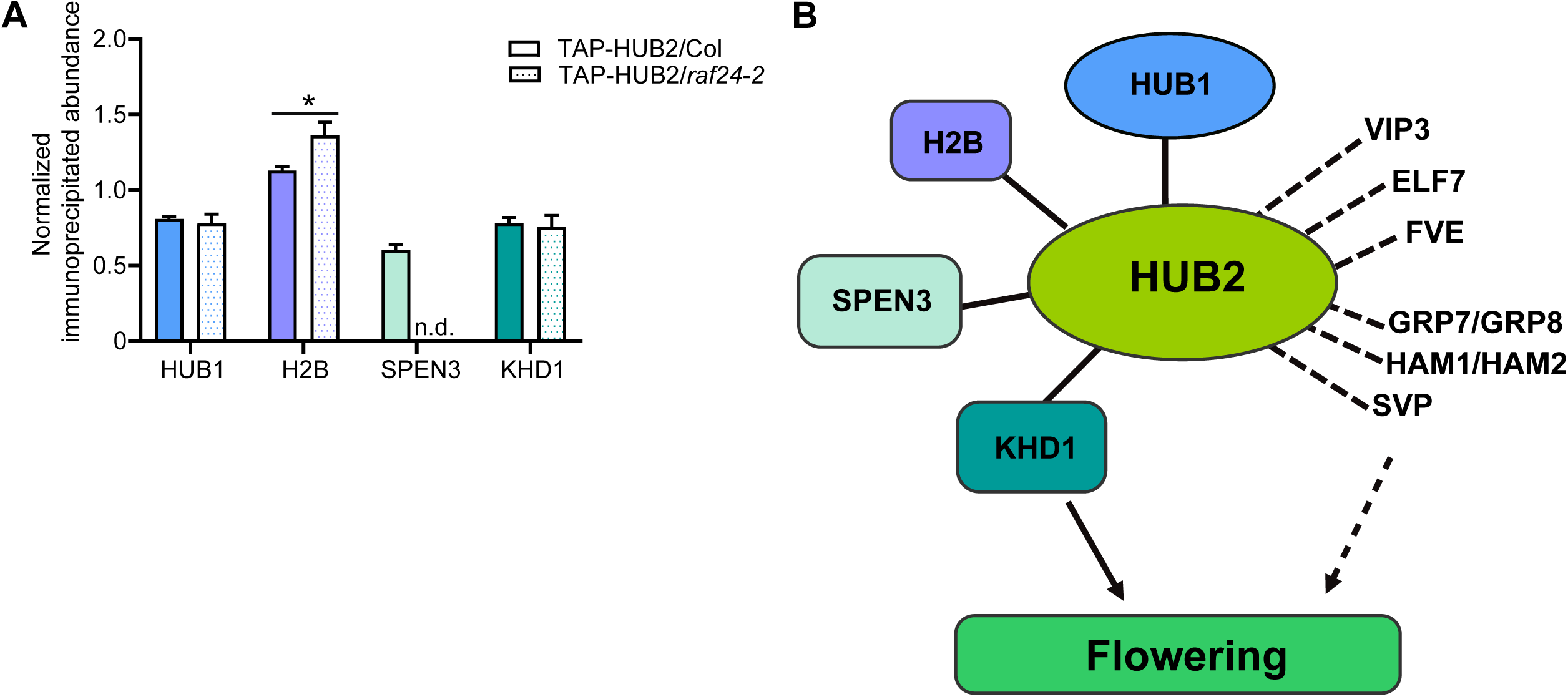
Loss-of-RAF24 impacts the interaction of HUB2 with H2B and SPEN3. (A) TAP-HUB2 pull-downs from 14d-old WT and *raf24-2* plants revealed altered association of known HUB2 interactors (n = 4). Quantification values were normalized to the abundance of TAP-HUB2 in either the TAP-HUB2/WT or TAP-HUB2/*raf24-2* backgrounds, respectively. (B) Schematic of quantified flowering-related interactors detected in TAP-HUB2 pull-down experiments (Supplemental Data 3). Known interactors were represented by colored bubbles with a black line. Novel flowering-related candidate HUB2-interactors were shown as bold text with a dash line.

Next, we queried our HUB2 interactome data for changes in the abundance of other known HUB2 interactors. Here we found a quantifiable decrease in the pulldown of the HUB2 interactor SPEN3 (AT1G27750), which is also known to affect flowering time (Woloszynska *et al*., 2019). Compared to WT Col-0, where a functional RAF24 is present, SPEN3 went undetected in TAP-HUB2 pulldowns from *raf24-2* plants, while equal amounts of the protein KHD1 (AT1G51580) (Woloszynska *et al*., 2019) was found in pulldowns from both WT Col-0 and *raf24-2* (Figure 6A) Supplemental Data 3). Further assessment of *SPEN3* and *KHD1* expression levels in both WT Col-0 and *raf24-2* by RT-qPCR found no change in their expression level (Supplemental Figure 6), confirming that absence of RAF24 affects the interaction between RAF24 and SPEN3, but not expression levels of *SPEN3*. Taken together, the protein-protein interaction strength of HUB2 and its known interactors, is impacted by the presence/absence of RAF24 and the resulting phosphorylation of HUB2.

Apart from these known flowering-related protein interactors of HUB2, we also found several characterized flowering regulators, including VERNALIZATION INDEPENDENCE 3 (VIP3; AT4G29830), HISTONE ACETYLTRANSFERASE OF THE MYST FAMILY 1/2 (HAM1/2; AT5G64610/ AT5G09740) and GLYCINE RICH PROTEIN 7/8 (GRP7/8; AT2G21660/AT4G39260), were pulled down by TAP-HUB2. (Figure 6B; Supplemental Data 3). Many candidate binding partners regulate flowering time in an FLC-dependent manner, but in different signaling pathways. VIP3 and EARLY FLOWERING 7 (ELF7; AT1G79730) being part of *Arabidopsis* relative of yeast RNA POLYMERASE II ASSOCIATED FACTOR 1 (PAF1) complex, modulate vernalization-responsive flowering time via FLC (Zhang *et al*., 2003; He *et al*., 2004; He, 2009; Cao *et al*., 2015), further supporting the previous hypothesis that HUB2 and PAF1 play roles in the same pathway (Wright *et al*., 2011). In addition, MULTICOPY SUPPRESSOR OF IRA1 4 (MSI4/FVE; AT2G19520) acts as a primary repressor of FLC in autonomous flowering signaling (Kim *et al*., 2004; Sun *et al*., 2021). Together, this suggests that in addition to affecting H2Bub1 levels, HUB2 may play additional regulatory roles in FLC expression by interacting with different binding partners upon upstream environmental cues. On the other hand, SHORT VEGETATIVE PHASE (SVP; AT2G22540), which serves as a transcriptional repressor of flowering in concert with FLC, is also found as a candidate interactor of HUB2 and has been found to be controlled by the 26S proteasome (Lee *et al*., 2013; Jin & Ahn, 2021). However, unlike the human HUB1/2 ubiquitin ligase orthologs RNF20/40, HUB2 has not been found to polyubiquitinate substrates for proteasomal degradation (Jeon *et al*., 2020). Therefore, future studies could investigate whether HUB2 polyubiquitinates SVP and if this affects its control over flowering initiation.

### SnRK2s interact with and phosphorylate HUB2 *in planta*

From our TAP-HUB2 pull-down data, we also found a number of protein kinases that possibly interact with HUB2 (Figure 7A). In particular, SnRK2s and CASEIN KINASE II (CK2), which have each been implicated in the regulation of flowering time (Figure 7A) (Ogiso *et al*., 2010; Mulekar & Huq, 2012; Wang *et al*., 2013). Interestingly, HUB2 was previously found to be a direct phosphorylation target of SnRK2.6 (Wang et al., 2020). In addition, previous studies showed that SnRK2.4 and SnRK2.6 are phosphorylated and activated by RAF24 (Lin et al., 2020; Soma et al., 2020; Fàbregas et al., 2020). Therefore, we hypothesized that RAF24 may impact HUB2 phosphorylation through RAF24-related SnRK2s. To test this, we first confirmed which SnRK2s interact with HUB2. Using a bimolecular fluorescence (BiFC) assay through transient co-expression of nYFP-HUB2 with each of the *Arabidopsis* SnRK2s, we find that fluorescence was only reconstituted with SnRK2.2-cYFP, SnRK2.4-cYFP or SnRK2.6-cYFP (Figure 7B; Supplemental Figure 7). These results suggest that SnRK2.2, SnRK2.4 and SnRK2.6 can interact with HUB2 *in vivo*, corroborating previous findings that SnRK2.4 and SnRK2.6 are directly phosphorylated by RAF24 (Lin et al., 2020; Soma et al., 2020). We next focused on the relationship between SnRK2.4 and SnRK2.6 with HUB2. First, we conducted split-luciferase complementation assays to further confirm interactions between HUB2-SnRK2.4 and -SnRK2.6. Here, we detect a significantly larger luciferase signal when co-expressing HUB2 with SnRK2.4 and SnRK2.6, versus SnRK2.8 (Figure 7C). Next, we tested if SnRK2.4 and SnRK2.6 phosphorylate HUB2 by conducting an *in vitro* ADP-Glo protein kinase activity assay. We found that HUB2 assays conducted with SnRK2.4-His_6_ or SnRK2.6-His_6_ displayed significantly higher relative light units (RLUs), implicating SnRK2.4 and SnRK2.6 in the phosphorylation of HUB2 *in vitro* (Figure 7D). Lastly, we examined if SnRK2.4 and SnRK2.6 phosphorylates HUB2 *in vivo*. To this end, we co-expressed the TAP-HUB2 and SnRK2s-YFP in *Nicotiana benthamiana*. Upon isolating TAP-HUB2, we found HUB2 to possess increased phosphorylation when co-expressed with SnRK2.4 or SnRK2.6, compared to that of SnRK2.8 (Figure 7E). Collectively, these results indicate that SnRK2.4, in addition to SnRK2.6, physically interact with, and phosphorylates, HUB2.

**Figure 7.**
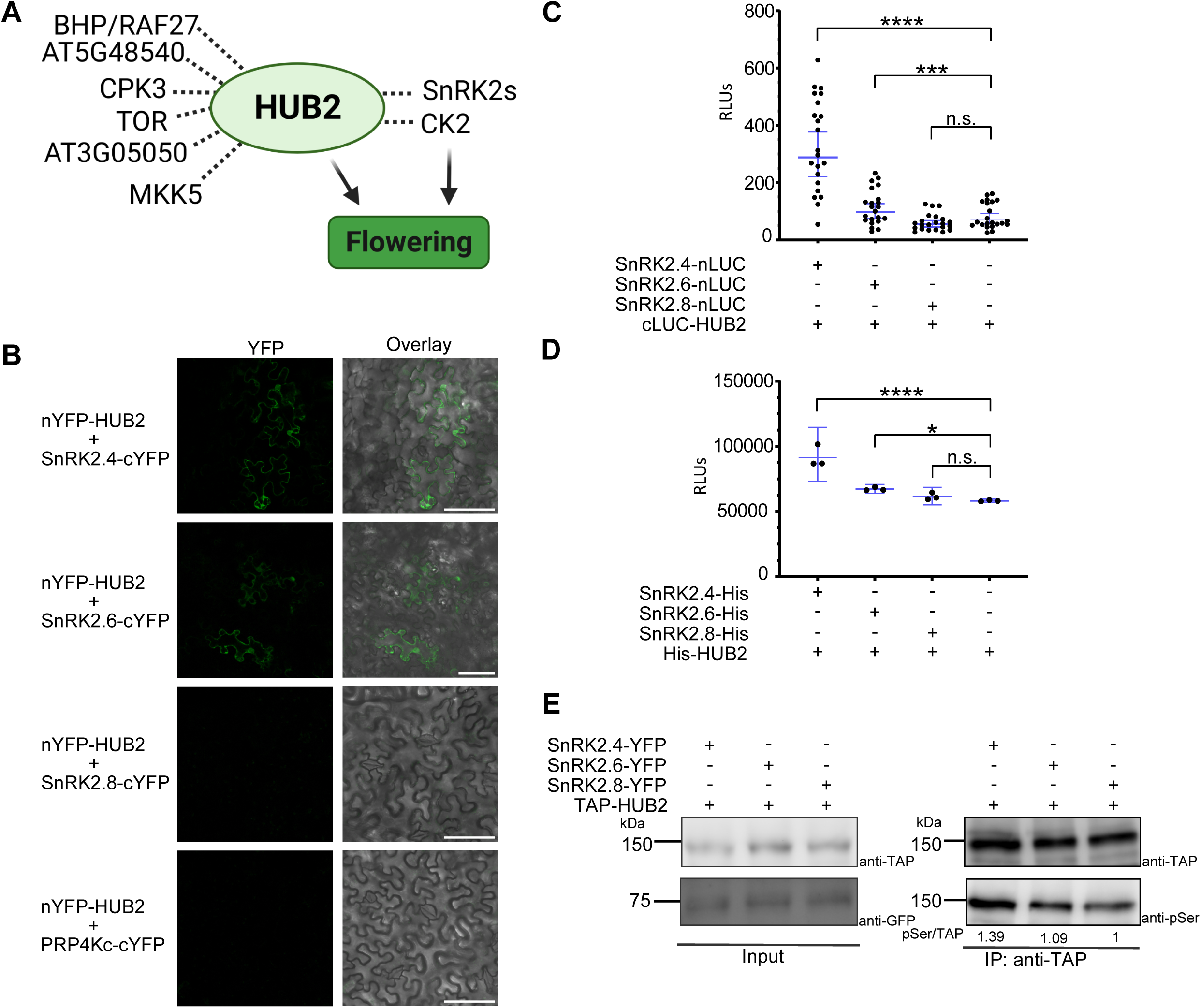
SnRK2.4 and SnRK2.6 interacts with and phosphorylates HUB2. (A) TAP-HUB2 pulldowns identified multiple protein kinases, including flowering-related SUCROSE NONFERMENTING 1-RELATED KINASE 2 (SnRK2) and CAESIN KINASE 2 (CK2) kinases. (B) Bi-molecular fluorescence complementation analysis validates interactions between HUB2 with SnRK2.4 and SnRK2.6 *in planta*. Co-infiltration of nYFP-HUB2 and SnRK2-cYFP constructs alongside PRP4Kc-cYFP (negative control) was performed using *N. benthamiana* leaves. Fluorescence signals were visualized with confocal microscopy 2 to 3d after infiltration (n > 10 images per set; Scale bars = 10 μm). (C) HUB2 interacts with SnRK2.4 and SnRK2.6, but not SnRK2.8 in luciferase complementation assays. Pairs of SnRK2s-nLUC and cLUC-HUB2 constructs were co-expressed using *N. benthamiana* leaves. After 3d incubation, luciferase intensity was quantified (n = 22 plants; Stars (*) denote one-way ANOVA *p-value* = 0.0001 (***) or *p-value* ≤ 0.0001 (****). (D) Phosphorylation of recombinant HUB2 by SnRK2.4 and SnRK2.6 *in vitro*. Relative HUB2 phosphorylation when mixed with recombinant SnRK2s or water (negative control). ADPglo luciferase signals were quantified after 3 h (n = 3; Stars denote one-way ANOVA *p-value* < 0.05 (*) and *p-value* ≤ 0.0001 (****)). (E) Phosphorylation of HUB2 by SnRK2.4 and SnRK2.6 *in vivo*. Co-expression of TAP-HUB2 and either YFP-tagged SnRK2.4, 2.6 or 2.8 was conducted through infiltrated *N. benthamiana* leaves. Immunoblotting analysis of each co-expression set was performed with anti-TAP, anti-GFP as well as anti-Ser IgG. 50 μg of input protein samples were loaded in each lane.

### Phosphorylation of HUB2 at S^314^ represses flowering and ligase activity

Lastly, to understand if phosphorylation of HUB2 at S^314^ translates to control of flowering time, we next created phospho-mimetic (HUB2^S314D^) and phospho-ablative (HUB2^S314A^) expressing transgenic *raf24-2* plants. We observed HA-HUB2^S314D^ expressing *raf24-2* plants to possess a significant delay in flowering time relative to both the HA-HUB2^S314A^ and HA-HUB2^S314S^ *raf24-2* plants (Figure 8A&B). The delayed flowering phenotype indicates that phosphorylation of HUB2 at S^314^ in the *raf24-2* background directly influences the flowering time of Arabidopsis. In addition, we observe higher H2Bub1 levels in HA-HUB2^S314A^ than HA-HUB2^S314D^ in the *raf24-2* background (Figure 8C), further confirming that phosphorylation of HUB2 at S^314^ suppresses its ligase activity.

**Figure 8.**
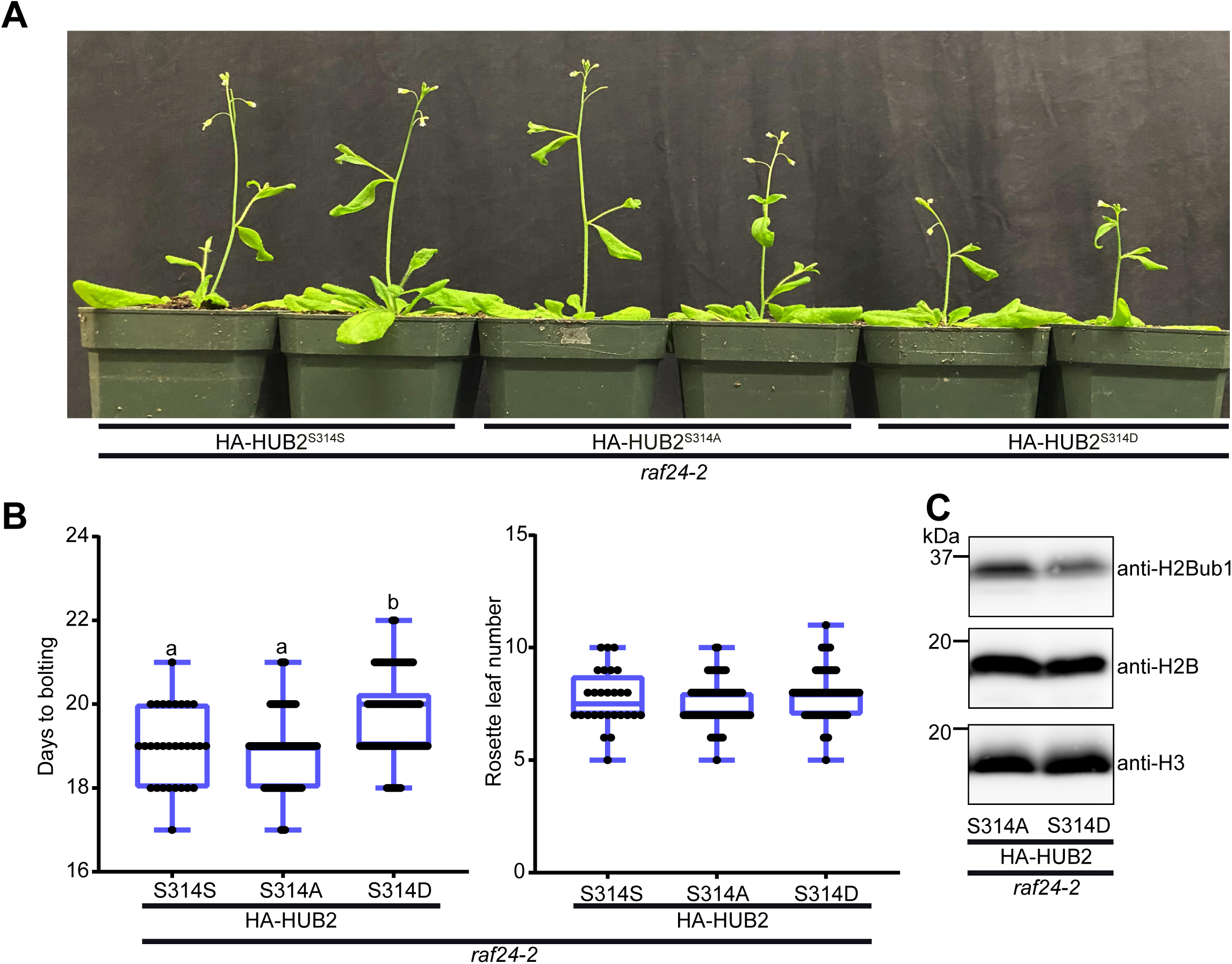
Phosphorylation of HUB2 at S^314^ controls flowering time. (A) and (B) Comparison of flowering time in *raf4-2* plants constitutively over-expressing HA-HUB2 with no (S314S), phospho-ablative (S314A) and phospho-mimetic (S314D) mutations (A). Flowering time was determined by days to bolting and rosette number at bolting (B). Letters indicate significant differences between genotypes (one-way ANOVA *p-value* < 0.05). (C) Immunoblot analysis of *raf24-2* plants constitutively over-expressing phospho-ablative (HA-HUB2^S314A^), phospho-mimetic (HA-HUB2^S314D^), which finds HA-HUB2^S314D^ maintains lower H2Bub1 levels. 50 μg of protein samples were loaded in each lane. Equal amounts of HA-HUB over-expression in the *raf24-2* background was observed across the lines depicted here (Supplemental Figure 7).

## DISCUSSION

Recent research has resolved roles for RAF-like MAPKKK kinases in abscisic acid (ABA) (Lin *et al*., 2021; Kamiyama *et al*., 2021), osmotic signaling (Lin *et al*., 2020; Soma *et al*., 2020), auxin signaling (Kuhn et al., 2024) and embryonic development (Wang *et al*., 2018b), with functions in flowering time remaining undefined. Here we identify a novel function for RAF24 in the control of flowering, which diverges from its shared roles with other B4 RAF MAPKKK kinases in osmotic stress responses where their phosphorylation of SnRK2 kinases facilitates down-stream signaling (Lin *et al*., 2020; Soma *et al*., 2020) and auxin signaling (Kuhn et al., 2024). Our identification of an independent role for RAF24 can be explained by its divergent phylogenetic relationship to its most closely related orthologs RAF18 and RAF20 and by their differential gene expression profiles, where we find RAF24 to be expressed in rosette leaves, versus a predominant root expression profile for RAF18 and RAF20 (MAPK Group, 2002; Soma *et al*., 2020). This suggests an intrinsic difference between the function of different B4 RAF MAPKKK protein kinases as it pertains to developmental regulation versus stress responses. In particular, in the early onset of abiotic stress, plants may employ several kinases to rapidly transduce stress signals, however, a single protein might be sufficient to induce certain changes in plant development.

### RAF24 sits high in flowering-time signaling, and may crosstalk with brassinosteroid and/or abiotic stress signaling

Here we find evidence that suggests RAF24 regulates flowering initiation through a number of pathways. First, phosphoproteomic analysis of the *raf24-2* mutant found several phosphoproteins related to flowering are potential RAF24 substrates (Table 1). Among these, VIP4 is responsible for vernalization-gated flowering time, while BES1 and FBH3 are involved in the photoperiod flowering regulation (Zhang & van Nocker, 2002; Ito *et al*., 2012; Wang *et al*., 2019)(Table 1). These results suggest RAF24 might gate signal transduction of multiple flowering pathways. In addition, RAF24 also may play roles in brassinosteroid (BR) signaling. Several BR-related phosphoproteins, BES1, BRI1 SUPPRESSOR 1 (BSU1)-LIKE 1 (BSL1/2; AT4G03080 / AT1G08420) and BES1/BZR1 HOMOLOG 4 (BEH4; AT1G78700), displayed significantly lower phosphopeptide abundance in *raf24-2* phosphoproteomic analysis (Table1). The potential involvement of RAF24 in BR signaling may explain why *raf24-2* mutants exhibited smaller rosette compared to WT, as BR is known to regulate cell expansion (Haubrick & Assmann, 2006; Zhiponova et al., 2013) (Supplemental Figure 3). In fact, BR signaling is also associated with flowering time (Li et al., 2010; Li et al., 2018). Therefore, RAF24 may mediate flowering via crosstalk with BR signaling by directly or indirectly modulating phosphorylation status of BR-related proteins. Furthermore, we find that loss of RAF24 leads to lower HUB2 phosphorylation at S^314^ through SnRK2.4 and SnRK2.6, which directly interact with, and phosphorylate, HUB2. Considering that RAF24 phosphorylates both SnRK2.4 and SnRK2.6, and that SnRK2.6 has been reported to control flowering time in plants (Wang *et al*., 2013; Chong *et al*., 2022), our results indicate that RAF24 phosphorylates SnRK2.4 and SnRK2.6 to impact HUB2 phosphorylation status and thereby fine tune flowering time. Furthermore, this RAF24-SnRK2s-HUB2 signaling module indicates RAF24 sits high in flowering initiation pathways. This differs from the flowering kinases SK12 and CPK32 described to date, which directly phosphorylate key flowering-time proteins, CO and FCA, respectively (Chen *et al*., 2020; Li *et al*., 2022). Recently, SnRK2.6 in tomato (*Solanum lycopersicum*) was reported to promote flowering time upon drought stress conditions (Chong *et al*., 2022). Since HUB2 has been related to abiotic stress tolerance (Zhou *et al*., 2017; Ma *et al*., 2019; Chen *et al*., 2019), it may be interesting to investigate whether RAF24-SnRK2s-HUB2 module mediates flowering time following exposure to abiotic stress stimulus. Therefore, further research can be conducted on roles of RAF24 in mediated flowering time under varying hormone or abiotic stress conditions.

### Appropriate flowering time requires H2Bub1 homeostasis, which depends on tightly regulated HUB2 ligase activity by phosphorylation

Similar to its human ortholog, we find HUB2 to also be a phosphoprotein in plants, with its phosphorylation status involved in controlling HUB2 functionality and maintaining appropriate H2Bub1 levels (Moyal *et al*., 2011). So far, H2Bub1 homeostasis has been linked to flowering time by multiple studies (Cao *et al*., 2008; Schmitz *et al*., 2009; Woloszynska *et al*., 2019), with *hub2* plants displaying lower H2Bub1 levels and an earlier flowering time (Cao *et al*., 2008). Conversely, loss of UBIQUITIN-SPECIFIC PROTEASE 26 (UBP26) results in an accumulation of H2Bub1, and also early flowering initiation (Schmitz et al., 2009). Therefore, we propose that precise phospho-mediated regulation of H2Bub1 levels is required in order to avoid premature flowering. Our results support this model, as loss of RAF24 leads to a significant increase in H2Bub1 levels, therefore accelerating flowering initiation. Similarly, in the *raf24-2* background, HA-HUB2^S314A^ had increased accumulation of H2Bub1 and accelerated flowering relative to HA-HUB2^S314D^, indicating that phosphorylation of HUB2 acts to suppress the overaccumulation of H2Bub1 to avoid premature flowering. However, HUB2^S314D^ did not completely rescue the early flowering phenotype compared to the *raf24* mutant, as an overall increase in HUB2 levels also initiates premature flowering (Woloszynska *et al*., 2019). However, HA-HUB2^S314D^ did complement the early flowering phenotype in *raf24* plants expressing comparable levels of HUB2 (HA-HUB2^S314S^).

To ensure H2Bub1 homeostasis, HUB2 ligase activity needs to be tightly regulated. In Arabidopsis, activity of ubiquitin ligases has been found to be modulated by protein kinases, such as SnRK2s (Ding *et al*., 2015a,b). For example, SnRK2.6 phosphorylates RING-type CHY ZINC-FINGER AND RING PROTEIN1 (CHYR1) to enhance its ligase activity in ABA and drought response signaling (Ding *et al*., 2015b). In addition, SnRK2.6 also phosphorylates PUB25/PUB26 in response to freezing treatment, therefore increasing its E3 ligase activity (Ding *et al*., 2015a). Likewise, here we show that SnRK2.6, as well as SnRK2.4, interacts with and phosphorylates HUB2. So far, H2Bub1 is the only characterized substrate of HUB2 (Liu et al., 2007). Therefore, it is reasonable to use changes in H2Bub1 levels to reflect the ligase activity of HUB2. Correspondingly, we observed higher H2Bub1 levels in the absence of RAF24, indicating that RAF24 functions to suppress HUB2 ligase activity. Interestingly, H2Bub1 levels are known to be influenced by osmotic stress in Arabidopsis (Chen *et al*., 2019), and by ABA treatment in rice (Ma *et al*., 2019); both biological responses related to RAF24 (Lin *et al*., 2020; Soma *et al*., 2020). Since SnRK2.4 and SnRK2.6 are also activated by osmotic stress and ABA, respectively, it is possible that changes of H2Bub1 levels in response to abiotic stress, are in part, controlled by different SnRK2s and their phosphorylation of HUB2.

In addition to exerting roles in flowering pathway, HUB2 is also known to regulate many other biological processes, but specific protein interactors of HUB2 in these biological contexts have not been determined. Our TAP pulldown results reveal an array of candidate HUB2 interactors involved in diverse signaling pathways linked to a wide range of biological processes (Supplemental Data 3). Beyond flowering, HUB2 also functions in plant innate immunity (Hu *et al*., 2014; Zou *et al*., 2014). Here, we find immunity-related proteins MODIFIER OF SNC1, 2 (MOS2; AT1G33520), MODIFIER OF SNC1,4 (MOS4; AT3G18165), PECTIN METHYLESTERASE 17 (PME17; AT2G45220) and RECOGNITION OF PERONOSPORA PARASITICA 13 (RPP13; AT3G46530) present as candidate binding partner for HUB2 (Bittner-Eddy *et al*., 2000; Rose *et al*., 2004; Zhang *et al*., 2005; Monaghan *et al*., 2009; Del Corpo *et al*., 2020). In addition, HUB2 affects the dynamics of microtubules (Hu *et al*., 2014), and here we find multiple microtubule-associated proteins MICROTUBULE-ASSOCIATED PROTEINS 65-1 (MAP65-1; AT5G55230) and MAP65-6 (AT2G01910) as candidate HUB2 interactors (Mao *et al*., 2005; Lucas *et al*., 2011). Moreover, RESPONSIVE TO DEHYDRATION 21A (RD21A; AT1G47128), SNW/SKI-INTERACTING PROTEIN (SKIP; AT1G77180) and ABA INSENSITIVE 8 (ABI8; AT3G08550) are also present as candidate HUB2 interactors, implicating HUB2 in abiotic stress resistance (Brocard-Gifford *et al*., 2004; Ma *et al*., 2019; Chen *et al*., 2019; Liu *et al*., 2020; Zhang *et al*., 2022), with RD21A having been reported to be polyubiquitinated (Kim & Kim, 2013). Thus, it is possible that HUB2 may also be involved in the polyubiquitination of RD21A for 26S proteasomal degradation. Lastly, H2Bub1 implicated in circadian regulation through candidate HUB2 interactors BASIC LEUCINE ZIPPER 63 (bZIP63; AT5G28770) and GRP7 (AT2G21660)(Heintzen *et al*., 1997; Staiger *et al*., 2003; Frank *et al*., 2018). Further studies based on such protein-protein interaction information could help advance our insights into HUB2’s diverse roles in regulating plant development and fitness.

In summary, we propose a model whereby RAF24 functions to repress flowering initiation by modulating the phosphorylation of HUB2 through SnRK2.4 and SnRK2.6 (Figure 9A). In the presence of RAF24, HUB2 phosphorylation is maintained, which in turn optimizes HUB2ub1 levels and maintains effective transcriptional activation of FLC (Figure 9A). Upon the loss of RAF24 HUB2 is no longer phosphorylated, resulting in an over-accumulation of H2Bub1, and reduced transcriptional activation of FLC (Figure 9A). In addition, we find that RAF24 acts upstream of a variety of diverse flowering time pathways, suggesting a wide-ranging role in fine-tuning floral development in response to different internal or external cues that goes beyond HUB2 (Figure 9B). Moreover, the diverse set of HUB2 candidate interactors found here indicates that HUB2 may function as part of a broad series of protein complexes that impact flowering initiation (Figure 9B), each of which warrants further investigation in future studies.

**Figure 9.**
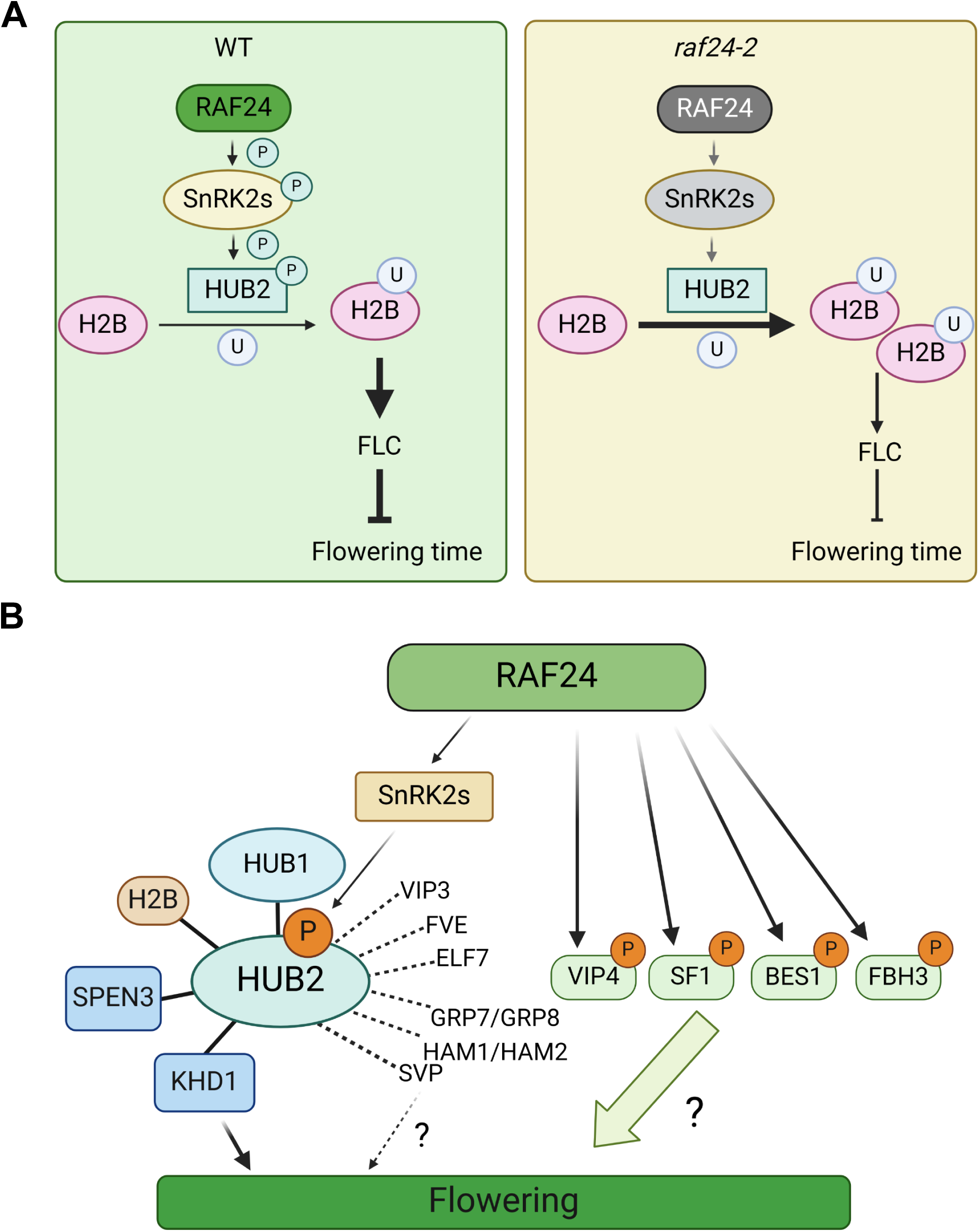
Roles of RAF24 in regulating HUB2 and flowering time. (A) In WT, the presence of RAF24 phosphorylates SnRK2s to induce phosphorylation of HUB2, ensuring appropriately tuned HUB2-ligase activity. This in turn optimizes HUB2ub1 levels to maintain proper transcriptional activation of FLC expression and flowering time (Figure 8A). Upon the loss of RAF24, SnRK2 fails to phosphorylate HUB2, causing an over-accumulation of H2Bub1 and reduced transcriptional activation of FLC expression. This results in earlier flowering (B). Both the phosphoproteomic and HUB2-TAP pulldowns find that RAF24 acts upstream of a wide variety of floral regulators, including BES1 and FBH3 as well as HUB2 to fine-tune flower time. Identification of both known HUB2 interactors (with straight line) and new, candidate HUB2 interactors (with a dash line) related to flowering time indicates that RAF24 possesses versatile functions / mechanisms in controlling flowering time.

## METHODS

### Plant Growth and Phenotyping Assays

#### Plant Materials and Growth Conditions

All Arabidopsis seeds used in this study were based on Col-0 ecotype. T-DNA insertion lines, *raf18-1* (SALK_143032), *raf20-1* (SALK_053369), *raf24-2* (SALK_107170), *raf24-4* (GABI_776D02), *raf24-5* (GABI_702G09), *hub2-1* (CS9779), *flc-6* (SALK_041126) as well as *ft-10* (CS9869) were obtained from The Arabidopsis Information Center and confirmed by genotyping evidence. The *raf24-2*, *hub2-1* were crossed to generate *raf24-2 hub2-1* double mutants. Similarly, *raf24-2*, *flc-6* and *ft-10* mutants were crossed to create *raf24-2 flc-6* and *raf24-2 ft-10* double mutants. All genotyping primers are listed in Supplemental Table 1. Arabidopsis seeds were stratified under 4°C and dark conditions for 3 days. Then all seeds were moved to a 16:8 or 12:12 light : dark photoperiod at 22 °C with a light intensity of 100 μE m−2 s−2 unless otherwise specified.

#### Leaf area measurement

Each chamber was equipped with a single-board Raspberry Pi 3 B+ computers and ArduCam Noir Cameras (OV5647 1080p) with a motorized IR-CUT filter and two infrared LEDs. Pictures of plants were taken every 15 min over seven days (between 14-d and 20-d post-seed imbibition). Using the open-source software package PlantCV (https://plantcv.readthedocs.io/), the areas of each plant in the picture collection were extracted and quantified as described (Gehan *et al*., 2017).

### Molecular Cloning and Transgenic Plant Generation

N-terminal TAP-HUB2G plants were created using the 35S::pKNGSrhino-HUB2G (TAP-HUB2) construct (Woloszynska *et al*., 2019). *Agrobacterium tumefaciens* GV3101 were propagated in 2ml liquid culture consisting of LB media containing 50 µg/µl Spectinomycin, 25 µg/µl Rifampicin and 40 µg/µl Gentamicin at 28°C 150 rpm for 48 h followed by additional 24 h sub-culturing at 1:20 dilution with the same antibiotic regiment. *Agrobacterium* was then pelleted and re-suspended in 40 ml 5% (w/v) sucrose, 10 ml LB and 3% (v/v) Silwet L-77 immediately prior to transfection. Floral dipping was performed as previously described (Clough & Bent, 1998). The transgenic seeds were selected on 0.5x MS addition with 50 µg/µl Kanamycin plates and screened under compound microscope (Olympus BX51 Fluorescence Microscope®) for fluorescence, prior to confirmation by western immunoblotting. For complementation line generation, the genomic RAF24 was amplified from cDNA library and then cloned into pEarlygate102 vectors. Following *Agrobacterium tumefaciens* GV3101 transformation, *raf24-2* plants were dippped to create RAF24-HA *raf24-2* complementation lines. RAF24 expression of complementation lines was verified by RT-qPCR. Subcellular localization assays utilized RAF24 and HUB2 cloned into pK7YWG2 and pEearlygate104 plasmids, respectively. For BiFC assays, the full-length coding sequence of each SnRK2 and HUB2 were cloned into pB5GWcY and pB5nYGW constructs, respectively. Lastly, HUB2 phospho-mimetic and phospho-ablative over-expressing lines were created using site-directed mutagenized 35S::pKNGSrhino-HUB2G (Supplemental Data 2), followed by insertion into the binary vector pEarlygate201. After transformation into GV3101, *raf24-2* plants were dipped to generate to HA-HUB2^S314^, HA-HUB2^S314A^, or HA-HUB2^S314D^ plant lines.

### Quantitative Phosphoproteome Analysis

Protein extraction, quantification, digestion and phosphopeptide enrichment were performed as previously described (Uhrig *et al*., 2019). Dissolved peptides were injected using an Easy-nLC 1000 system (Thermo Scientific) and separated on 50 cm ES803 Easy-Spray PepMap C18 Column (Thermo Scientific). Peptides were eluted using the following gradient of solvent B (0.1% (v/v) FA in 80% (v/v) ACN): 0–50 min, 0–25% B; 50–60 min, 25–32% B; 60–70 min, 32–98% B, at a flow rate of 300 nL/min at 50°C. Mass spectra were acquired using an Orbitrap Hybrid mass spectrometer (Thermo Scientific) operated in data-dependent acquisition mode as previously described (Uhrig *et al*., 2019). Raw data were analyzed using MaxQuant software version 2.0.3.0 (https://www.maxquant.org/; (Cox & Mann, 2008; Tyanova *et al*., 2016)) using default settings, which included: fixed modifications (cysteine carbamidomethylation; C), variable modifications (serine, threonine, and tyrosine phosphorylation; pSTY & methionine oxidation; M_ox_) and peptide, protein and PSM false discovery rate of 1%. Data outputs were processed using Perseus software version 1.6.15.0 (https://www.maxquant.org/; (Tyanova *et al*., 2016) and subsequently filtered for significantly changing phosphopeptides (students t-test; *p-value* < 0.05) with a data completeness of ∼65% (5 / 8 samples) and a PTM-site probability score ≥ 0.75.

### Affinity Purification Mass Spectrometry (AP-MS)

Positive TAP-HUB2 in Col-0 (TAP-HUB2/Col-0) and *raf24*-2 (TAP-HUB2/*raf24-2*) seedlings were grown for 14 d and harvested. Proteins were extracted using 50 mM HEPES-KOH (pH7.5), 100 mM NaCl, 0.1% (v/v) NP-40, 50 mM NaPPi, 1 mM Na_3_VO_4_, 2 mM PMSF, 1x Roche cOmplete™ EDTA-free protease tablet and 1x Roche PhosSTOP™ phosphatase inhibitor cocktail tablet (Roche). Extracts were clarified at 20,000 x g for 20 minutes at 4°C and then measured by Bradford Assay (Bradford, 1976). Extracts were then incubated with custom magnetic IgG-coupled beads (Cytiva) end-over-end for 1 h at 4°C to bind TAP-HUB2. IgG-coupled beads were then washed 3x for 5 min with 0.1% (v/v) NP-40 and 4 washes with no NP-40, followed by elution using 100 mM Glycine pH 3.0 neutralized with 1M Tris-HCl. TAP-HUB2 pull-downs were then reduced (10mM Dithiothreitol; 45min @ 37°C), alkylated (55 mM Iodoacetamide; 1h @ 23°C) and digested in solution with 1:100 trypsin (V5113; Promega). Peptides were acidified to 0.5% (v/v) trifluoroacetic acid (TFA; Sigma), dried, dissolved in 3% (v/v) ACN / 0.1% (v/v) TFA and desalted as previously described (Uhrig *et al*., 2019). Differences in the Col-0 and raf24-2 HUB2-protein interactome were quantified by BoxCarDIA LC-MS/MS analysis as previously described (Mehta et al., 2022). Acquired data was then analyzed using Spectronaut v14 (Biognosys AG) using default settings. Candidate HUB2 interactor abundance was then normalized against HUB2 levels to obtain relative fold-changes in abundance between each genotypic background.

### Co-localization and Bimolecular fluorescence complementation (BiFC) analysis

RAF24-YFP, mCherry-HUB2, nYFP-HUB2, SnRK2s-cYFP as well as PRP4Kc-cYFP were transformed into *Agrobacterium* through electroporation. Positive colonies were selected, cultured and harvested in infiltration buffer (10 mM MES pH 5.6, 10 mM MgCl_2_, 100 µM acetosyringone) as described (Schütze *et al*., 2009). Cell suspensions were diluted to OD = 0.8 to 1.0 and incubated in dark for at least 2 hours at room temperature before injecting 4-week-old *N. benthamiana* plants. Infected plants were grown for 2 - 3d, with infiltrated sections analyzed with a Leica SP8 STED or ZEISS LSM 710 facilities.

### Split-Luciferase Complementation Assay

Split-Luciferase complementation assays were conducted based on previously reported (Chen et al., 2008). *A. tumefaciens* GV3101 harboring cLUC-HUB2 (pGWB-CLUC) and SnRK2s-nLUC (pGWB-NLUC) were infiltrated into *N. benthamiana* as described above. Three days after infiltration, leaf discs were collected and placed into 96-well plate. Then 1 mM D-luciferin (Cayman, 25836) was applied into each well and incubated for 10-15 minutes in dark before luminescence measurement with the SPARK® multimode microplate reader (TECAN).

### Recombinant Protein Expression And in Vitro ADP-Glo Kinase Assays

His_6_-HUB2 (pDEST17) and SnRK2s-His_6_ (pET-DEST42) were used for protein expression. Briefly, BL21-CodonPlus cells containing His_6_-HUB2 and SnRK2.4-His_6_ were grown at 4 °C for 48 hours, while SnRK2.6-His_6_ and SnRK2.8-His_6_ were grown at room temperature for 24 h. Lysis was performed by French Press, with whole cell extracts cleared by centrifuged at 20000 x g at 4 °C degree for 30 minutes. Supernatants were retained and passed by miracloth, then incubated with His_6_-tag Purification Resin (Roche, 05893682001) at 4 °C for 1 hour. This was followed by three washes buffer A (50 mM HEPES pH 7.5, 500 mM NaCl, 1% (v/v) Tween-20) and three washes buffer B (50 mM HEPES pH 7.5, 150 mM NaCl. Proteins were eluted (50 mM HEPES, 150 mM NaCl, 300 mM imidazole) and concentrated using 10000 MWCO concentrator spin-columns (Millipore). Final protein concentrations were measured using Bradford’s reagent (Thermo Fisher).

Phosphorylation of HUB2 by SnRK2s was examined with ADP-Glo Protein Kinase Activity Assay Kit (Promega, V9101) with slight modifications (Kathania *et al*., 2022). 2 µg of His_6_-HUB2 was mixed with or without 1 ug of SnRK2s-His_6_ in kinase reaction buffer (50 mM Tris-HCl pH 7.5, 20 mM MgCl_2_, 100 µM ATP, 2 mM DTT and 1 mM EGTA) and incubated at 30 °C for 1 hour and at room temperature for 2 hours. Assays were stopped by adding ADP-Glo reagents and incubation for 40 minutes in the dark. Kinase activity detection reagent was then added and kept at room temperature for 30 minutes, luminescence signals were quantified with a SPARK® microplate reader (TECAN).

### *In Vivo* Phosphorylation Assays

*Agrobacterium* harboring TAP-HUB2G and SnRK2s-GFP (pB7YWG2) were co-expressed in *N. benthamiana.* Leaves were collected and ground using a mortar and pestle after 3 d, then extracted with protein extraction buffer (50 mM HEPES pH 7.5, 150 mM NaCl, 1 mM EDTA, 10% (v/v) glycerol, 2% (w/v) PVPP, 100 mM DTT, 0.1% (v/v) NP-40, 1 mM PMSF, 1x Roche cOmplete™ EDTA-free protease tablet and 1x Roche PhosSTOP™ phosphatase inhibitor cocktail tablet (Roche)) end-over-end for 30 minutes. Protein extracts were then incubated with custom magnetic IgG-coupled beads (Cytiva) for 1 hour at 4 °C and washed five times (50 mM HEPES pH 7.5, 100 mM NaCl). Beads were then mixed with 60 µL Tris-HCl pH 8.0 and 20 µL of 4x Laemmli Sample Buffer (BIO-RAD) for 10 minutes at 95 °C. Standard procedures were performed to visualize the TAP-HUB2, SnRK2s-GFP as well as phosphorylated HUB2 proteins by using anti-PAP (1:1000, SIGMA, P1291), anti-GFP (1:2000, ChromoTek, PABG1) and anti-phosphoserine antibodies (1:250, abcam, ab9332), respectively.

### Western Immunoblot Analysis

#### Protein Extraction and Preparation

10-day-old Arabidopsis seedlings were homogenized (2010 Geno/Grinder®) and then incubated with extraction buffer (4% (w/v) SDS, 50 mM Tris-HCl pH 8.0) for 30 minutes in an end-to-end rotator at 4 °C. Next, supernatants were retained after a 30-minute spin down with 16,000 x g at 4 °C. Protein estimation was conducted with Pierce BCA Protein Assay kit (Thermo Scientific). Finally, protein samples were mixed with 4x Laemmli Sample Buffer (BIO-RAD) for 5 minutes at 95 °C or 10 minutes at 65 °C.

#### Western Immunoblotting

Samples were loaded onto 7.5% or 15% SDS-PAGE gels and electro-transferred with Trans-Blot® SD Semi-Dry Transfer Cell (BIO-RAD) at 18 V for 45 minutes. After blocking 1 hour with 5% (w/v) non-fat milk, PVDF membranes were incubated overnight with anti-H2B (1:2500, abcam, ab1790), anti-H2Bub1 (1:1000, Cell Signaling, #5546), anti-GFP (1:2000, ChromoTek, PABG1), anti-HA(1:1000, Invitrogen, 26183), anti-H3 (1:2000, Invitrogen, PA5-16183) or anti-PAP (1:1000, SIGMA, P1291) followed by a 2-hour incubation with secondary antibodies, anti-rabbit (1:10000, Invitrogen, G-21234) or anti-mouse (1:5000, Invitrogen, G-21040).

### Quantitative PCR Analysis

#### RNA extractions

RNA was extracted from 4 biological replicate pools of 2-week-old Arabidopsis seedling. 100 mg of seedling tissue was harvested and ground using Geno/Grinder® for 30 seconds at 1200 rpm. Total RNA was extracted using a modified TRIzol based protocol (Chomczynski & Sacchi, 1987). 1 ml of TRI Reagent® (Sigma-Aldrich, T9424) was added per 100 mg of tissue and incubated for 10 minutes at room temperature. Extracellular material was removed by centrifuging at 13000 x g for 10 minutes. Supernatant was acquired and combined with 200 µl of chloroform. Mixture was then inverted variously for 15 seconds and then left at RT for 3 minutes. Phase separation was achieved by centrifuging at 13000 x g for 15 minutes at 4 °C. The aqueous phase was transferred to new tubes. RNA was precipitated by adding 500 µl of 100% (v/v) isopropanol and then left to incubate for 10 min at RT. Sample was centrifuged at 13000 x g for 15 minutes at 4 °C to acquire RNA pellet. Supernatant was removed and the pellet was washed with 1 ml of 75% (v/v) ethanol. Sample was centrifuged once again at 7500 x g for 5 minutes at 4 °C. Pellet was dried and subsequently resuspended in nuclease-free water. Total RNA was quantified by NanoDrop ND 1000 spectrophotometer (Thermo Scientific). Quality of RNA was assessed by visualizing 28S and 18S bands on a 1% (w/v) agarose gel stained with SYBRsafe (Thermo Fisher Scientific). Genomic DNA was removed using DNAse I kit (Thermo Fisher) followed by reverse transcription using RevertAid kit (ThermoFisher Scientific Inc) with 2 ug of starting RNA.

#### Transcript quantification

RT-qPCR was performed with 3 technical replicates per biological replicate using a proprietary mix developed and distributed by the Molecular Biology Service Unit (MBSU), in the Department of Biological Science at the University of Alberta. The SYBR mix contains Tris (pH 8.3), KCl, MgCl_2_, Glycerol, Tween 20, DMSO, dNTPs, ROX as a normalizing dye, SYBR Green (Molecular Probes, Life Technologies) as the detection dye, and an antibody inhibited Taq polymerase. A QuantStudio (TM) 6 Flex System was used, with UBC21 (AT5G25760) deployed as the reference for normalization. Transcript abundance was measured using delta CT method. Primer sequences are listed in Supplemental Data 2.

## ACKNOWLEDGEMENTS

The authors would like to thank Dr. Mieke Van Lijsebettens (Center for Plant Systems Biology, Belgium) for providing the 35S::pKNGSrhino-HUB2G construct, to acknowledge Dr. Hugo Zheng and Dr. Jiaqi Sun (McGill University) for sharing BiFC vectors and 35S::mCherry vectors, as well as thank Dr. Gitta Coaker (University of California, Davis) and Dr. Sonia Gazzarrini (University of Toronto) for providing SnRK2s clones. Further thanks go to Jack Moore of the Alberta Mass Spectrometry and Proteomics facility for mass spectrometry assistance.

## DATA AVAILABILITY

All raw proteomics data can be found in PRoteomics IDEntification database (PRIDE; https://www.ebi.ac.uk/pride/) PXD038688.

*Username:*

*Password: Teitvbdy*

## AUTHOR CONTRIBUTIONS

QL conducted phenotypic analysis and characterized phospho-mutants, in addition to performing all WB, confocal microscopy, phosphorylation assays as well as designing experiments. LW generated all seed sources and performed phenotypic analysis, TAP-pulldown experiments, proteomic experimentation and qPCR analyses unless otherwise stated. LEG performed RAF24 complementation line screening, as well as RT-qPCR and flowering assays for complementation lines. MR performed all COOLAIR qPCR analysis. MT performed protein purification and enzyme assays. DM assisted with experimental design and analysis. SS performed phenotypic analyses relating to plant surface area measurements and processing with PlantCV. RGU conceived of the study, while QL and RGU wrote the manuscript with editorial input from DM.

## SUPPLEMENTAL FIGURES

**Supplemental Figure 1.** Distinct phopsho-motifs of B4 like RAFs are phosphorylated in response to diel or osmotic stress changes. Relative abundance and retrieved from published phosphoproteomic analysis.

**Supplemental Figure 2. Loss-of-RAF24 induces premature flowering.** (A) and (B) Early flowering time of *raf24* mutants but not *raf18* and *raf20* mutants. 56-day-old plants were photographed under 12:12 photoperiodic conditions and flowering time scored by days to bolting and rosette leaf number at bolting. Letters indicate significant differences between genotypes (one-way ANOVA *p-value < 0.05*). (C) Relative expression analysis of *RAF24* in each *raf24* mutant. Transcript levels of *RAF24* were quantified using 14d-old *raf24-2*, *raf24-4* and *raf24-5* mutants with P1 and P2 primer pairs amplifying the 5’ and 3’ends of RAF24, respectively (Supplemental Data 2). Data were normalized to UBC21 expression levels and then relative to *RAF24* transcriptional levels in WT background. (D) Relative expression analysis of *RAF24* in two complementation lines of *RAF24-HA raf24-2* in comparison to Col-0 and *raf24-2*. Transcript levels of *RAF24* were quantified with P1 primer pairs amplifying the 5’ end of *RAF24* (Supplemental Data 2). Data were normalized to UBC21 expression levels.

**Supplemental Figure 3. Loss-of-RAF24 displays a smaller leaf area.** (A) and (B) The leaf area of 29-days post-imbibition WT and *raf24-2* plants was measured as described (see materials and methods), with *raf24-2* plants displaying reduced leaf area. Plants were grown and photographed under either 16:8 (upper panel) or 12:12 (bottom panel) photoperiodic conditions.

**Supplemental Figure 4. Localization of RAF24-YFP in *N. benthamiana* leaves.** Fluorescence signal of RAF24-YFP was detected by confocal microscopy 3d after infiltration. Scale bars = 20 μm.

**Supplemental Figure 5. Immunoblot analysis of H2Bub1 levels across *hub2* and *raf24-2/hub2* plants.** Immunoblot analysis was performed using 10-day-old plants. 50 μg of protein samples were loaded in each lane. Abundance of H3 proteins serves as the loading control.

**Supplemental Figure 6. Quantitative PCR analysis of known HUB2 interactors in WT and *raf24-2* plants.** Transcript quantification was performed using 14d-old WT and *raf24-2* plants. Data indicates that no statistically relevant change in transcript levels were found between WT and *raf24-2* genotypes. Student’s t-test *p-value* ≤ *0.05*.

**Supplemental Figure 7. HUB2 specifically interacts with SnRK2.2, SnRK2.4 and SnRK2.6.**

*N. benthamiana* leaves were co-infiltrated with nYFP-HUB2 and each cYFP-SnRK2s as per the materials and methods. Infiltration of nYFP-HUB2 alone served as a negative control along with PRP4Kc-cYFP (Figure 7). Fluorescence signals were visualized with confocal microscopy 3d after injection. n > 10 images were acquired for each set. Scale bars = 20 μm.

**Supplemental Figure 8. Immunoblotting of HA-HUB2 proteins levels when expressed in the *raf24-2* background.** 10d-old plants were harvested and analyzed by immunoblot. Ponceau staining serves as a loading control. 50 μg of protein samples were loaded in each lane. Equal amounts of HUB2 over-expression is observed across plants expressing plants expressing either HA-HUB2^S314^, HA-HUB2^S314A^, or HA-HUB2^S314D^.

## SUPPLEMENTAL DATA

**Supplemental Data 1: Phosphoproteome data**

**Supplemental Data 2: Primer sets used in the study**

**Supplemental Data 3: HUB2-TAP pulldown data**

**Supplemental Data 4:SnRK2 subclasses**

## Notes

### Competing Interest Statement

The authors have declared no competing interest.

### Summary of Updates

Figures were updated with additional experimentation

