## Supplementary figures and images for "B4 Raf-like MAPKKK RAF24 regulates *Arabidopsis thaliana* flowering time through HISTONE MONO-UBIQUITINATION 2"

### Supplemental Figure 1

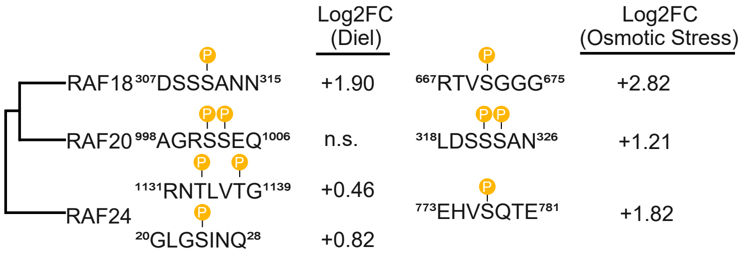

### Supplemental Figure 2

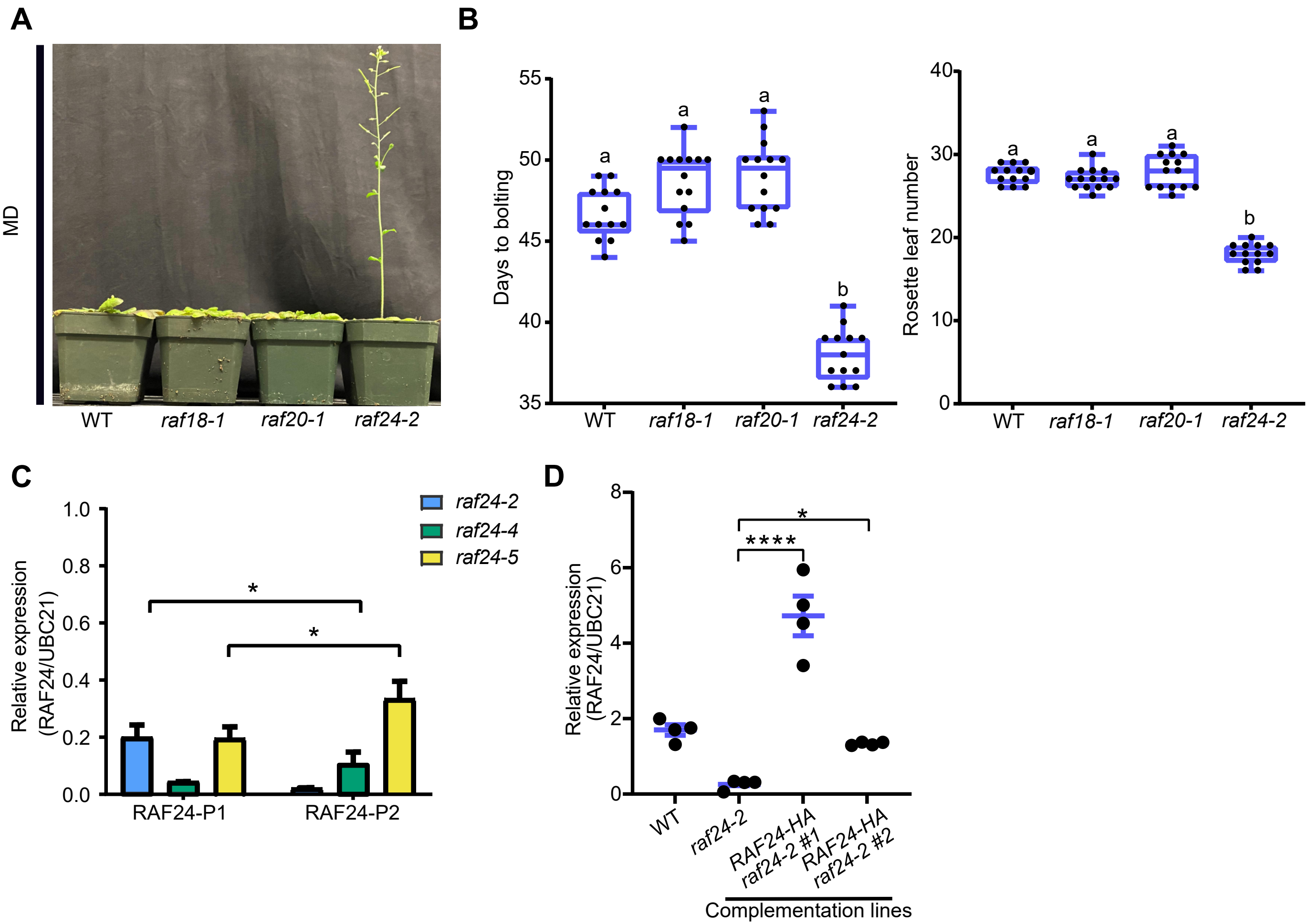

### Supplemental Figure 3

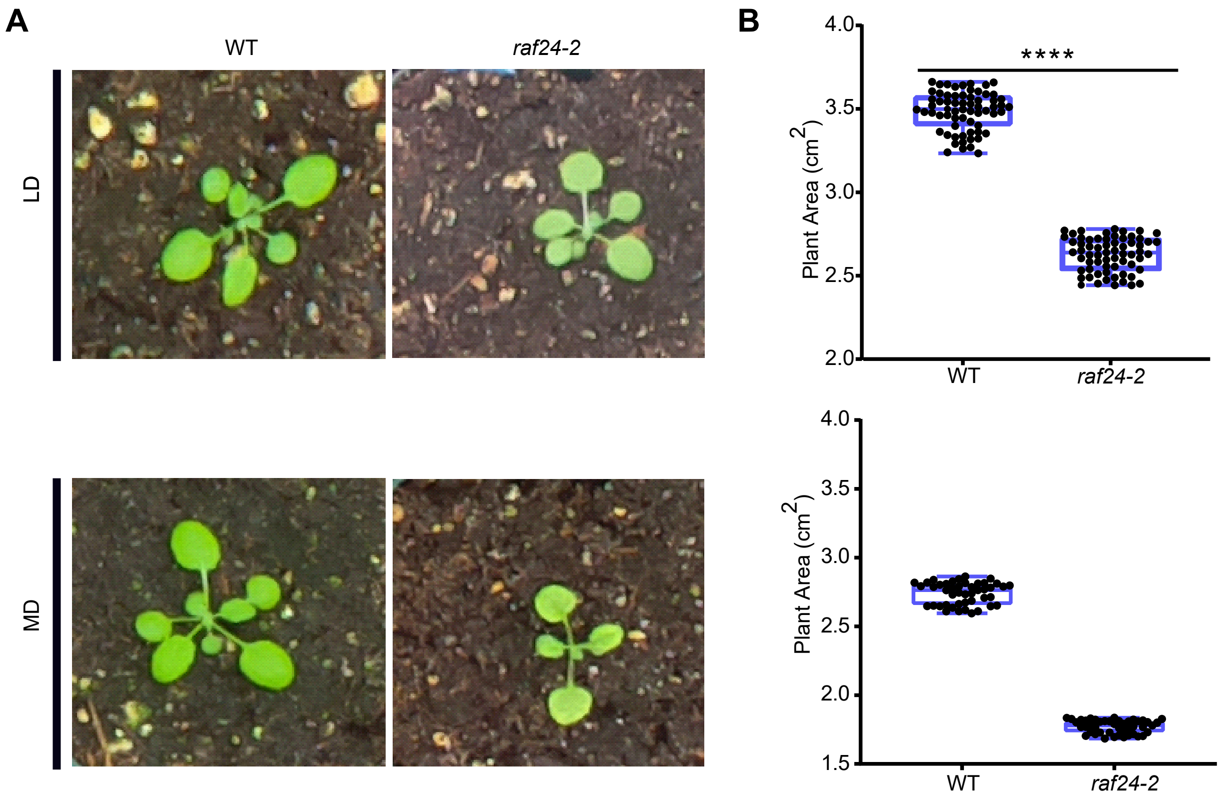

### Supplemental Figure 4

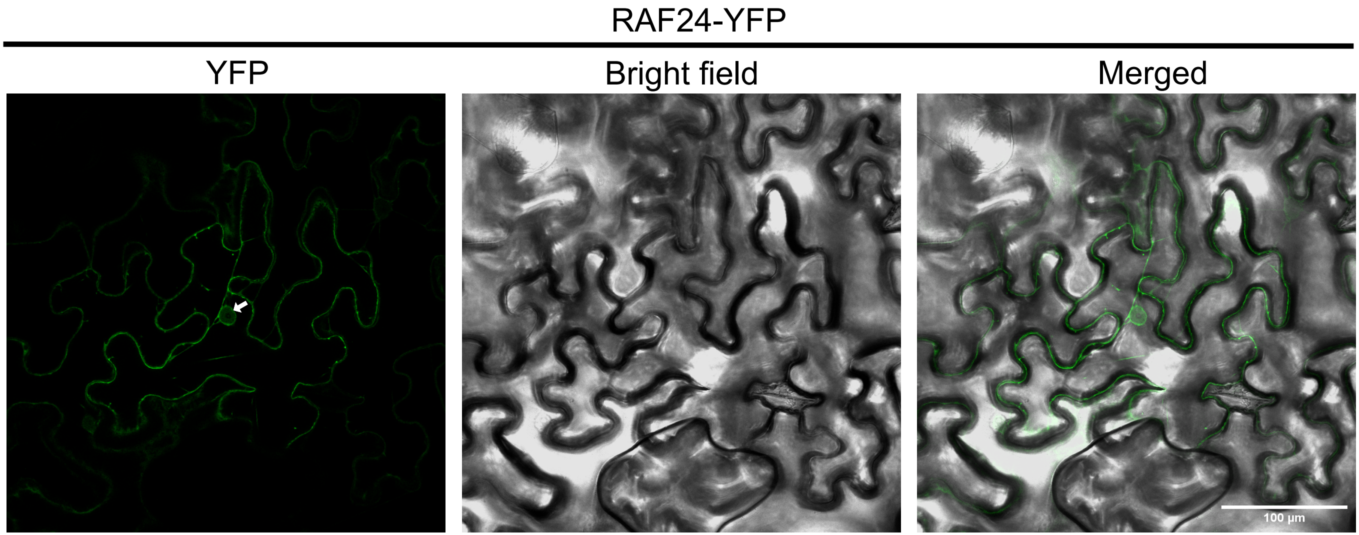

### Supplemental Figure 5

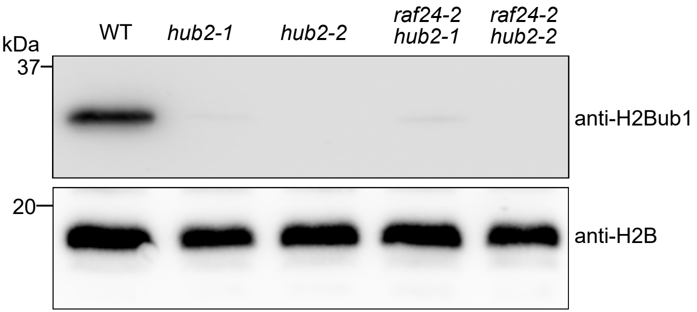

### Supplemental Figure 6

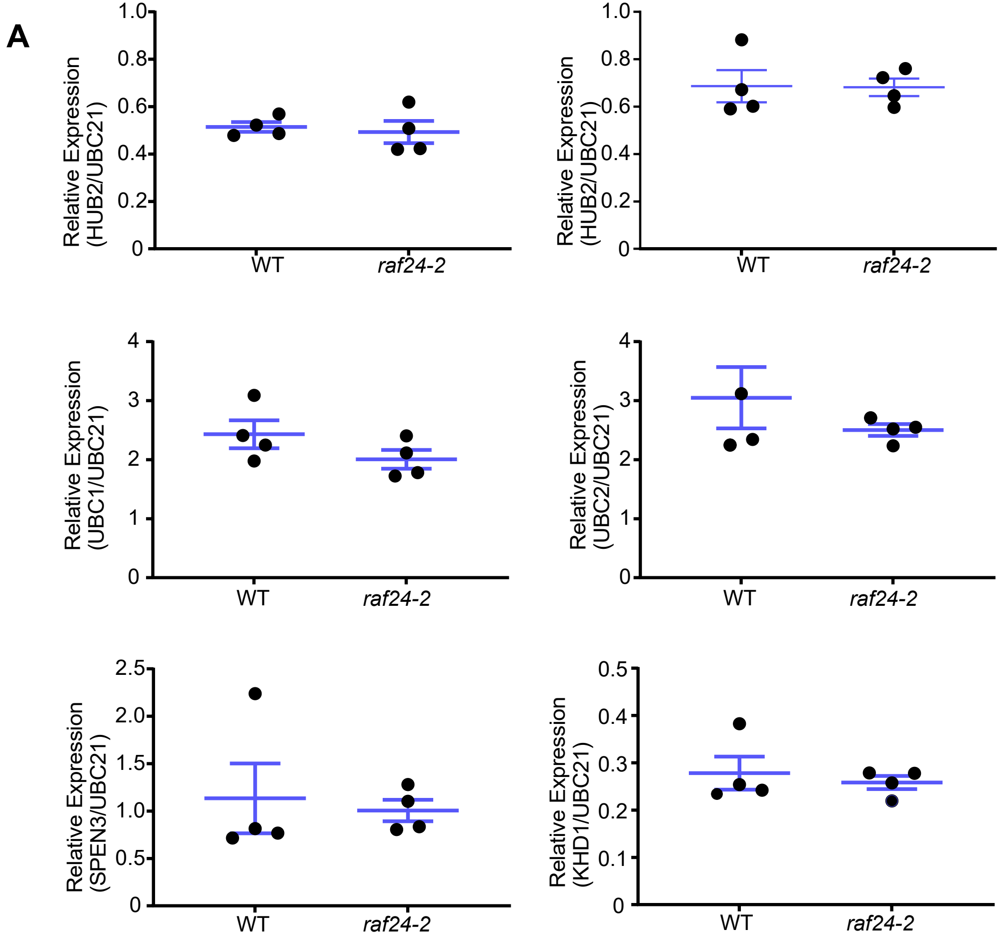

### Supplemental Figure 7

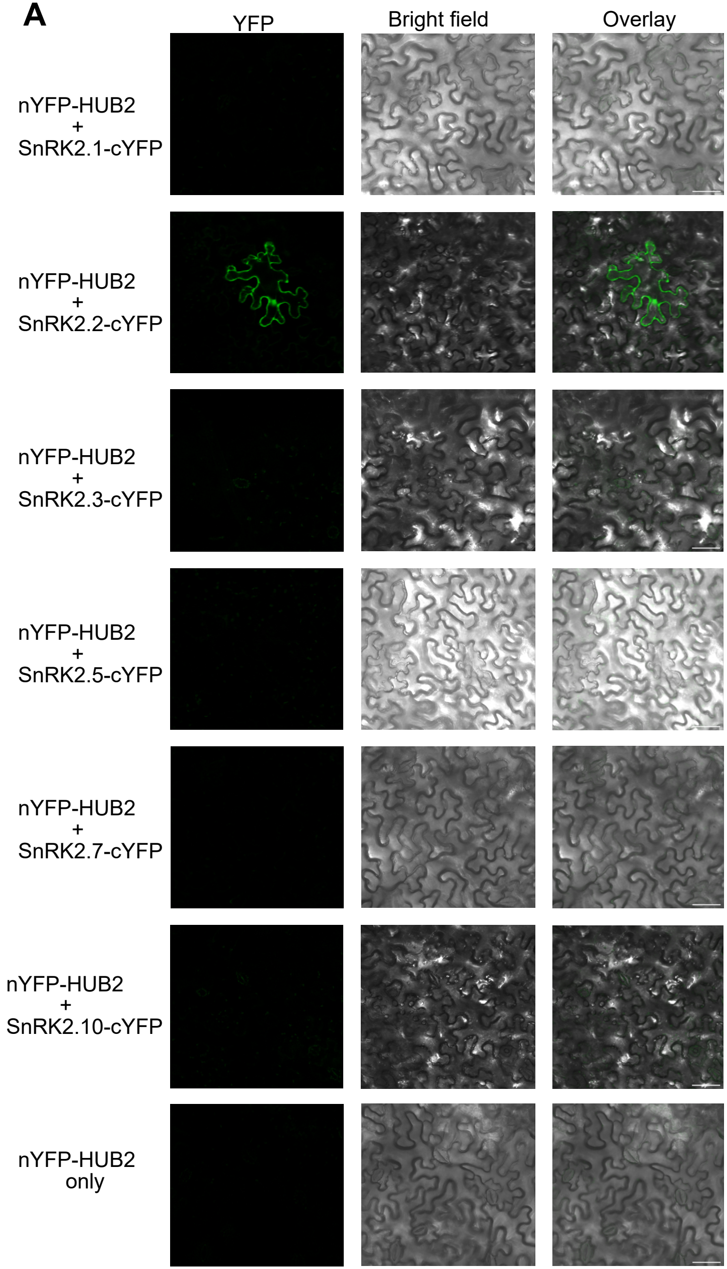

### Supplemental Figure 8

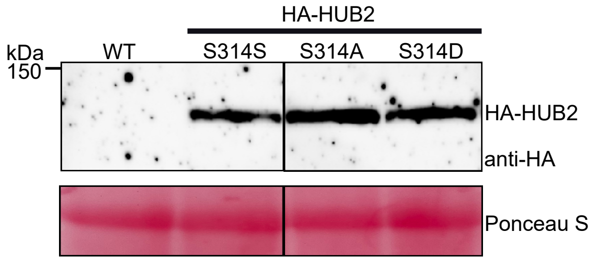
